# There is no single functional atlas even for a single individual: Parcellation of the human brain is state dependent

**DOI:** 10.1101/431833

**Authors:** Mehraveh Salehi, Abigail S. Greene, Amin Karbasi, Xilin Shen, Dustin Scheinost, R.Todd Constable

## Abstract

The goal of human brain mapping has long been to delineate the functional subunits in the brain and elucidate the functional role of each of these brain regions. Recent work has focused on whole-brain parcellation of functional Magnetic Resonance Imaging (fMRI) data to identify these subunits and create a functional atlas. Functional connectivity approaches to understand the brain at the network level require such an atlas to assess connections between parcels and extract network properties. While no single functional atlas has emerged as the dominant atlas to date, there remains an underlying assumption that such an atlas exists. Using fMRI data from a highly sampled subject as well as two independent replication data sets, we demonstrate that functional parcellations based on fMRI connectivity data reconfigure substantially and in a meaningful manner, according to brain state.

## Introduction

Human neuroscience research has long endeavored to assign specific functions to clearly demarcated brain regions. Neuroimaging has offered deep insights into functional specialization in the human brain, and has permitted the localization of functional regions. Brain mapping approaches have been focused on assigning specific roles to such regions and through this approach much has been learned about the functional organization of the brain. Recently, there has been significant interest in whole-brain parcellation approaches for deriving brain atlases ^1^. The parcellation problem itself is of great interest, and whole-brain parcellation is particularly relevant for defining the human connectome, which characterizes the interactions between brain regions. These developments have fueled a growing interest in functional connectivity-based analyses. There has been no consensus to date, however, on how to define the underlying atlas that best reflects the brain’s functional organization.

The widely used Brodmann areas^2^, defined by cytoarchitectural boundaries, were among the earliest attempts to subdivide the brain into functionally meaningful units, and there have been numerous, varied approaches to generate such atlases since^3-8^. Neuroimaging-based parcellation approaches are attractive because they allow whole-brain parcellations in individuals^9-12^ or groups of subjects^13-19^ (for a review see Eickhoff et al.^20^). Most recent neuroimaging-based parcellation algorithms have been based entirely on functional connectivity data^13-16^ or combinations of anatomical and functional data^17^,^18^. In addition, meta-analytic databases such as BrainMap^21^ and NeuroSynth^22^ have attempted to collate information from thousands of studies to provide behavioral context on any region of the brain, and clustering methods have been developed to translate these findings into homogeneous regions^23^. Yet all of these commonly adopted parcellations, whether at the individual or group level, define a single functional atlas with the underlying assumption that parcels are homogenous in function and invariant in size, shape or position regardless of brain state.

In this work, we provide evidence that there is not a single functional parcellation atlas but rather that the flexible brain reconfigures these functional parcels depending upon what it is doing. While there are many timescales on which brain states can be measured and many ways in which they can be defined (see Discussion), here we use tasks to elicit discrete, distinct brain states, and demonstrate that parcel boundaries change across task-induced states, yet are reliably reproducible within a state. Further, we show that the particular configuration of the parcels provides meaningful information on brain state: that is, a measure as coarse as parcel size for a given atlas can significantly predict the task condition under which the data were acquired, as well as the within-condition task performance. Using a single, highly sampled subject, where we know there are no anatomic differences across conditions or sessions, and wherein one would expect the parcels to be consistent from session to session, we demonstrate that the parcels are indeed consistent for a given condition, but reproducibly reconfigure across conditions, even when starting with the same initial atlas each time. These results hold across two additional independent data sets, both within and across subjects, and at various parcellation spatial resolutions, suggesting that they are the result, not simply of individual idiosyncrasies in functional brain organization or of systematically varying noise in boundary estimates, but rather of robust, generalizable, state-dependent reorganization of functional areas. This suggests that a single functional parcellation of the human brain is neither attainable nor desirable, it emphasizes the importance of considering brain state when drawing functional boundaries on the brain, and offers functional parcellations as a tool to study state-dependent changes in functional organization of the human brain. These findings provide another dimension with which to understand the organization of the flexible brain under changing brain state or cognitive conditions.

## Results

Three independent data sets are used to support these findings. In the first, 30 sessions of fMRI data were obtained from a single subject; each session was approximately 60 minutes long and included 6 task conditions (*n*-back^24^, gradual-onset continuous performance task [gradCPT]^25-27^, stop-signal [SST]^28^, card guessing^29^, Reading the Mind in the Eyes^30^, and movie watching) and 2 rest conditions. Data were acquired from a single subject to eliminate the potential confound of inter-individual variations in anatomy, which would contribute to the variance in parcel boundaries^31^. The high number of fMRI sessions acquired in a single subject allows us to demonstrate consistent within-condition parcellations across sessions, while also demonstrating significantly different cross-condition parcellations, both within and across sessions. Since a single subject was used, the null hypothesis would be consistent parcellations across all sessions and conditions. A second, independent data set was used from Midnight Scan Club (MSC)^32^ to replicate these findings and demonstrate their generalizability. This publicly available data set consists of a set of 8 individuals, scanned under both resting-state and task conditions 10 times (10 sessions) each. Finally, rather than measuring consistency within a subject across sessions, we used the Human Connectome Project (HCP)^33^ data (n=514) to demonstrate that even when collapsing across subjects (rather than sessions), we observe that different conditions lead to reproducibly different functional parcellations.

In the first data set, which we refer to as Yale data, we applied an extended version of our recently developed individualized parcellation algorithm^11,34^ to each functional run, generating one parcellation atlas for each condition and each scanning session (8 conditions × 30 sessions = 240 atlases total). A key factor in this algorithm is that each parcellation begins with an atlas obtained from a separate group of subjects and finds an exemplar time-course for each parcel and then grows the parcels (see Methods and Figure 7 for a detailed explanation of the algorithm). This exemplar-based approach has many advantages but the most important advantage for this work is that correspondence of parcels from different parcellations is maintained, making for straightforward comparisons of the resulting atlases. Most other parcellation approaches could yield completely different atlases, and in that case, it can be difficult to match parcels for quantitative comparison. However, to rule out the possibility that the parcellation algorithm choice affected presented results, we replicated the main findings with two other parcellation algorithms: (i) a modification of our exemplar-based method with fixed exemplars across sessions and conditions, and (ii) Wang’s iterative parcellation^35^ (see Methods, Figure S4 and Figure S5 for details).

**Figure 7.**
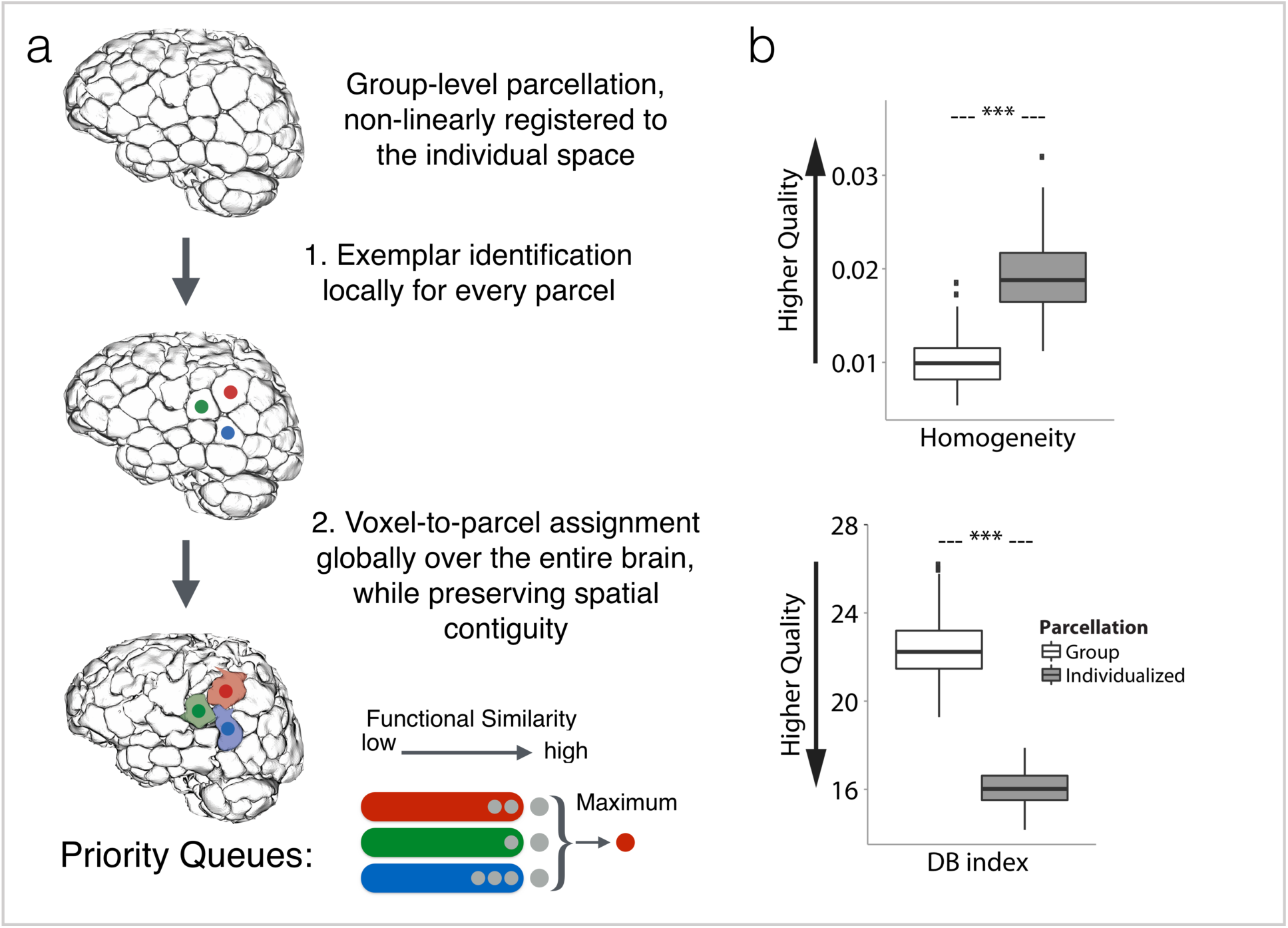
Individualized state-specific parcellation pipeline and evaluation. a) Three-step parcellation pipeline: Step 1: Starting from data in common space, a group-level functional atlas is first applied to the individual subject. Step 2: For every group-defined parcel in the individual brain, an exemplar is identified by maximizing a monotone nonnegative submodular function; here, three exemplars are shown in red, green, and blue. Step 3: Every voxel in the individual brain is assigned to the functionally closest exemplar while taking the spatial contiguity of the parcel into account. The spatial contiguity is assured by utilizing priority queues. Every exemplar *i* is assigned a priority queue (denoted as *q*_*i*_), here depicted as red, green, and blue queues which correspond to red, green, and blue exemplars, respectively. Initially, all the queues are empty. In the first round, spatial neighbors of exemplar *i* are pushed into *q*_*i*_. The voxels in each queue are sorted according to their functional distance to the corresponding exemplar such that the voxel with minimum functional distance (maximum similarity) is in front. Next, the front voxel in each *q*_*i*_ is considered as a potential candidate for being assigned the label *i*. Among all these candidates, the one with minimum distance to its corresponding exemplar is selected and assigned the exemplar’s label; here the candidate voxel from the ‘red’ queue is selected and labeled ‘red’. Next, this voxel is popped out of the queue, and all of its spatial neighbors are pushed into the same queue. The algorithm continues until all the voxels are assigned a label. Note that at every step of the algorithm the labeled voxel is ensured to be spatially connected to its exemplar (either directly or through other previously labeled voxels). b) Quantitative results for assessing the quality of parcellation. Homogeneity (top) and DB index (bottom) comparison between the individualized, state-specific parcellations and the initial group-level parcellation, represented as box plots with the central mark indicating the median, and the bottom and top edges of the box indicating the 25^th^ and 75^th^ percentiles, respectively. The whiskers extend to the most extreme data points not considered outliers, and the outliers are plotted individually. *** p <2.2e-16, two-tailed Mann-Whitney test.

By taking the relative majority vote over all sessions of each condition, we generated condition-specific parcellations. Figure 1a visualizes these parcellations using a force-directed graph, with edge weights indicating the similarity between parcellations, measured by r_Hamming_, which estimates the percentage of voxels with similar parcel assignment. Parcels are colored by the magnitude of reconfiguration in the given condition relative to Rest 1, where reconfiguration is defined as the percentage of voxels that change their parcel assignment. Figure 1a demonstrates that parcels with high reconfiguration are broadly distributed and condition specific. Increased similarity of reconfiguration maps is observed (Figure 1a) among condition-specific parcellations with increased pairwise similarity (Figure 1b). Consistent with our expectations, Rest 1 and Rest 2 parcellations are highly similar to each other, while the parcellation for the movie-watching condition is the most distinct (Figure 1). For visualization and interpretation purposes, we created a video of these state-specific parcellations (Video S1) as well as an interactive brain visualization platform (see http://htmlpreview.github.io/?https://github.com/YaleMRRC/Node-Parcellation/blob/master/Parcellation_visualization.html).

**Figure 1.**
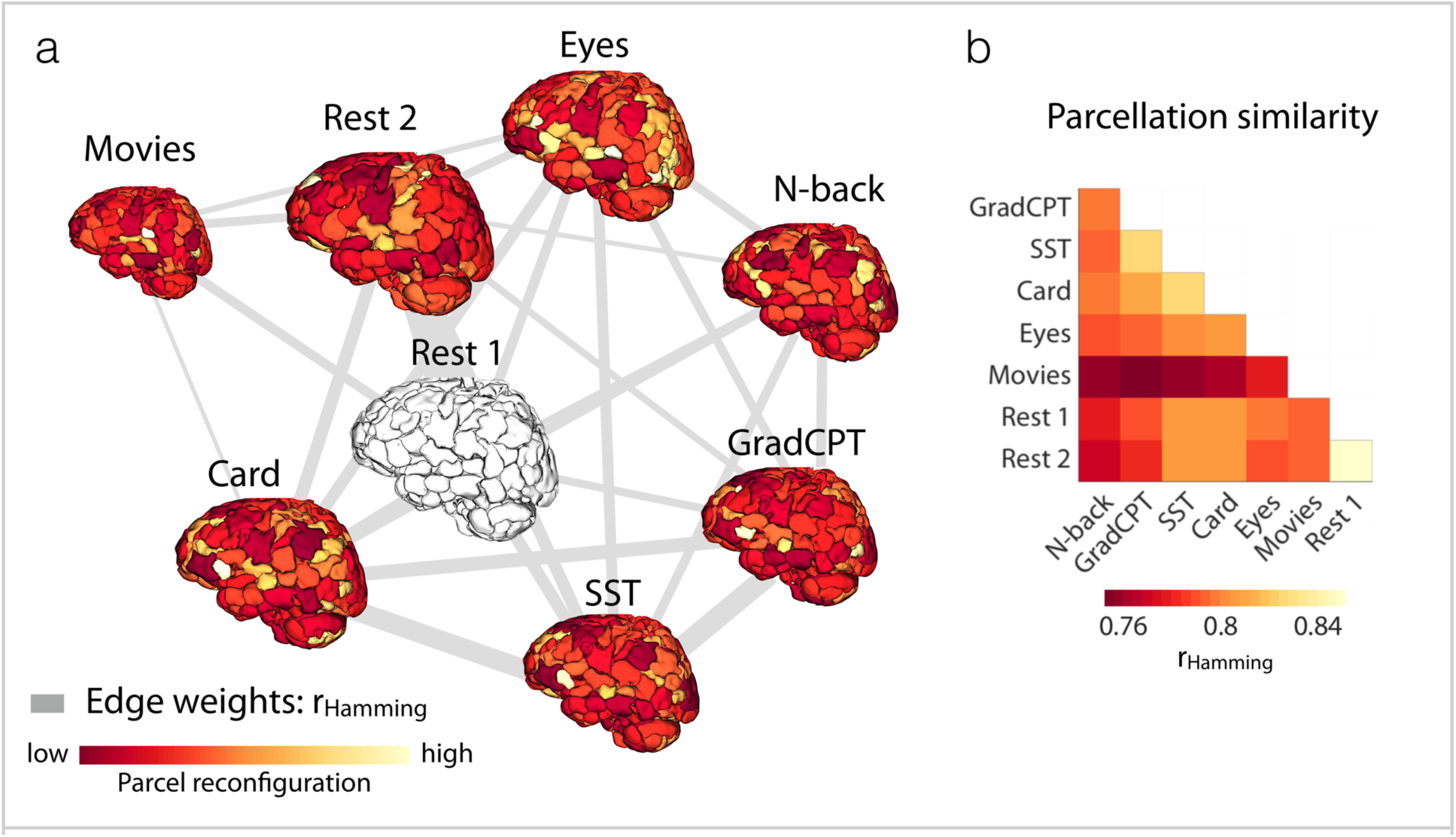
Visualization of the condition-specific functional atlases. (a) Condition-specific functional atlases are visualized in a force-directed graph, with edge weights indicating the similarity between parcellations, measured by r_Hamming_ = 1 − normalized Hamming distance. Force-directed graphs attempt to visually organize networks such that the energy of the graph as a whole is minimized. This is accomplished by assigning both repulsive and attractive forces to each pair of nodes such that the nodes with stronger interconnections are displayed closer to each other and the ones with weaker connections are more distant. Brain size is proportional to the graph theory measure degree. Edge thickness is proportional to the edge weights. Parcels are colored by the magnitude of reconfiguration in the given condition relative to Rest 1. (b) Cross-condition parcellation similarity measured by r_Hamming_.

To quantitatively examine these parcellation changes across conditions, we applied a statistical voting-based ensemble method, randomly dividing all sessions for a given condition into two equal-sized groups, obtaining the relative majority vote over each group, and computing the similarity between pairs of parcellations. This was done both within and across functional conditions. We repeated this analysis 1000 times, generating 1000 pairwise similarity matrices (see Methods for details). Parcellation similarity was considered at two scales: at the fine scale, we assessed the fraction of voxels that retained their parcel assignment (Figure 2a, d); at the coarse scale, we computed the rank correlation between parcel-size vectors (Figure 2b, e) for all condition pairs. The results demonstrate that parcellations are significantly more similar within each functional condition than across different conditions (K-S test; p<0.001). This finding also holds true when parcel size is considered as a means to summarize atlas boundaries (K-S test; p<0.001). The primary finding is that parcellations are highly reproducible within a functional condition, and rearrange significantly between conditions. These changes are reflected even in a coarse-scale feature of spatial topography, i.e. parcel-size.

**Figure 2.**
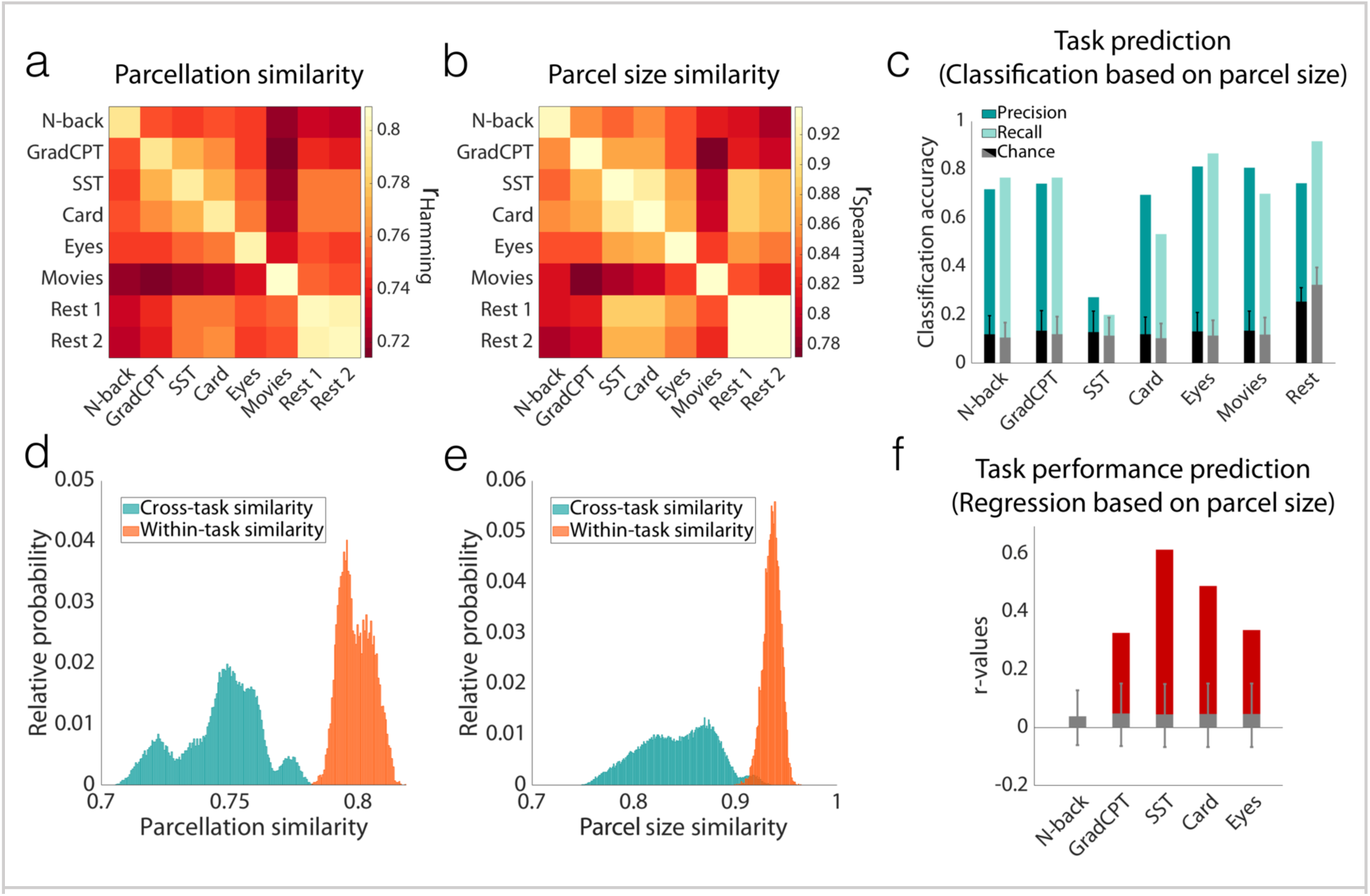
Parcel definitions change with task condition; Yale single-subject data. a) Pairwise parcellation similarity was calculated within and across functional conditions (8 conditions, N=30 sessions), using voting-based ensemble analysis with 1000 iterations. The matrix represents the average over all iterations. Similarity was assessed by r_Hamming_ = 1 – normalized Hamming distance. b) The same analysis as (a) was performed, this time using rank correlation of parcel-size vectors (r_Spearman_) as a proxy for parcellation similarities. c) The bars reflect the accuracy of predicting the functional condition using a leave-one-out cross-validated GBM classifier with parcel sizes as features. The predictive power is measured by precision (dark cyan) and recall (light cyan) values for each condition. The precision (black) and recall (gray) values of 1000 null models are also reported (error bars represent ±s.d.). d) A histogram of the parcellation similarities for all 1000 iterations is depicted for within-condition (diagonal elements in [a]) and cross-condition (off-diagonal elements in [a]) comparisons. Rest 1 and Rest 2 are grouped into one condition. The within- and cross-distributions are significantly different (K-S test; p<0.001). e) A histogram of the parcel size similarities for all 1000 iterations is depicted for within-condition (diagonal elements in [b]) and cross-condition (off-diagonal elements in [b]) comparisons. Again, the two distributions are significantly different (K-S test; p<0.001). f) The bars report the accuracy of predicting task performance using a leave-one-out cross-validated linear regression model with parcel sizes as features. Predictive power is measured by square root of coefficient of determination (r-value). The r-values of 1000 null models (for permutation testing) are also reported (error bars represent ±s.d.).

Next, we demonstrate that the observed task-induced parcellation reconfiguration is consistent across sessions and specific to each condition. To this end, we built a cross-validated predictive model that predicts the functional condition under which novel runs were acquired based solely on the parcel size vector. Prediction accuracies—measured as precision and recall for each condition—were significantly higher than random for all conditions (mean accuracy = 71%; Figure 2c). That a measure of spatial topography as coarse as parcel size can significantly predict which task a subject is performing suggests that functional brain atlases reconfigure with task-induced brain state in a consistent manner, forming a generalizable and robust signature of brain organization during that condition. Successful prediction also demonstrates that cross-condition reconfigurations are not driven by noise.

While the overall stability of the functional atlases within each condition was high, subtle variations were observed across different sessions of the same condition. To examine if parcel reconfigurations contain meaningful session-specific information, we attempted to predict task performance using parcel size as the model feature. Task performance was used as a proxy for level of engagement in the task, reflecting more fine-grained variations in brain state than cannot be captured by simply considering task as a state (see Methods for details of the behavioral measures that were used for each task). To this end, a linear regression model was employed in a leave-one-out cross-validated framework (see Methods for details), for each task independently. We used coefficient of variation (r-value) to assess the predictive power of the model. Task performance was successfully predicted from parcel sizes (Figure 2e), indicating that reconfigurations from session-to-session within a task condition reflect meaningful changes in brain function. The success of these predictions again rules out the possibility that even session-to-session reconfigurations are driven by noise.

Our final analysis aims to demonstrate that the state-evoked parcel reconfigurations are not driven by voxels with low certainty in their parcel assignment. To this end, we built upon our voting-based ensemble analysis and quantified voxel uncertainty as the proportion of times across 1000 iterations that a voxel was assigned to different parcels between the two groups. Figure 3a shows that the majority of the voxels have less than 10% uncertainty in their parcel assignment, suggesting that the parcellations are reliable within a functional condition. Figure 3b visualizes the spatial distribution of voxel uncertainty in the brain. From this visualization, it is clear that the shape of the white areas in each parcel, representing voxels with low uncertainty (uncertainty < 10%), changes substantially between conditions, suggesting that the state-evoked reconfigurations are not simply driven by noisy boundaries, but that stable voxels form parcels of different shapes.

**Figure 3.**
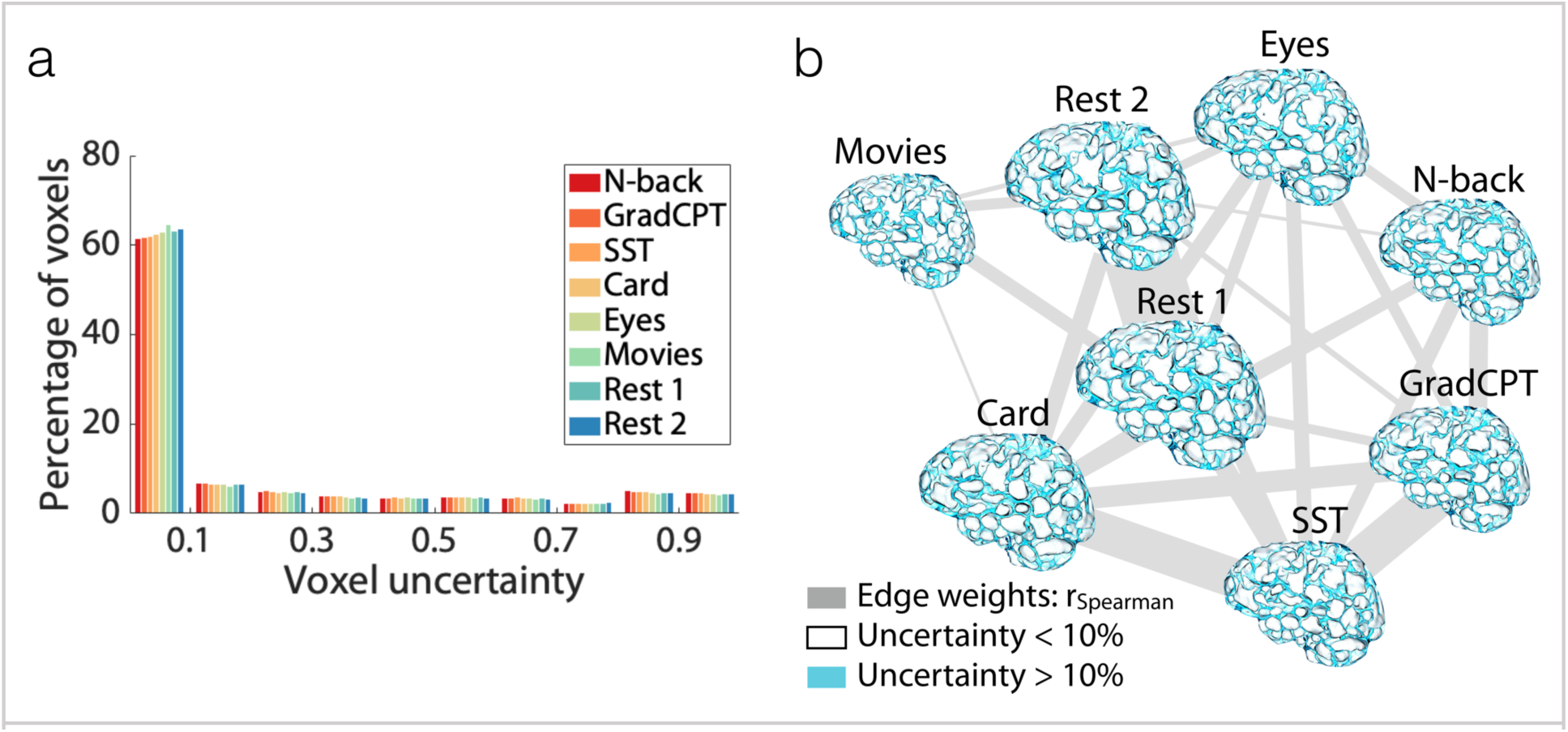
Voxel uncertainty analysis. Voxel uncertainty was computed for each functional condition as the proportion of times across 1000 iterations of the voting-based ensemble analysis that the voxel was assigned to different parcels between group 1 and group 2 (see Methods). a) The histogram of voxel uncertainty for each functional condition. b) The spatial distribution of voxel uncertainty on the brain, visualized in a force-directed graph with edge weights indicating the similarity between voxel uncertainty distributions across conditions, measured by r_Spearman_. Brain size is proportional to the graph theory measure degree. Edge thickness is proportional to the edge weights. Voxels with a low uncertainty (uncertainty < 10%) are shown in white, and the parcels defined by these clearly change shape between conditions. The rest of the voxels (uncertainty > 10%) are shown in blue.

### Replication of the state-evoked atlas reconfiguration across data sets

We replicated our main findings by leveraging a publicly available data set: Midnight Scan Club (MSC)^32^. The MSC data are particularly well suited for this analysis, as they include task-based and resting-state fMRI data from 10 individuals, each of whom was scanned 10 times. Each scan session included seven runs of tasks (semantic-coherence [2 runs], memory [scenes, faces, and words; each 1 run], and motor [2 runs] tasks) and 1 resting-state run (eyes open). Two individuals were excluded from the analyses, one (MSC10) due to missing data and the other (MSC08) due to excessive head motion (see Methods for details), leaving 8 individuals (4 females; age = 24 – 34) for analysis.

For each individual, we repeated the voting-based ensemble analysis described above and calculated the similarity between each task-(or rest-) based pair of parcellations. Figure 4 demonstrates the pairwise parcellation similarities, averaged over individuals (Figure 4a,d: fine-scale similarity; Figure 4b,e: coarse-scale similarity). Figure S1 shows the similarity matrices of each individual separately. Confirming the finding from the single-subject data described above, we observed that functional atlases are significantly more similar within condition than across conditions (K-S test, p<0.001). Similarly, parcel sizes are significantly more similar within a condition (K-S test, p<0.001). Predictive modeling based on parcel size could significantly predict condition for a novel run (mean accuracy = 66%; Figure 4c) and generalized across individuals. That all the predictions were successful, and consistent across individuals, is a strong indication of the significance of functional parcel reconfigurations associated with different tasks. The findings from this independent data set replicate the single-subject findings.

**Figure 4.**
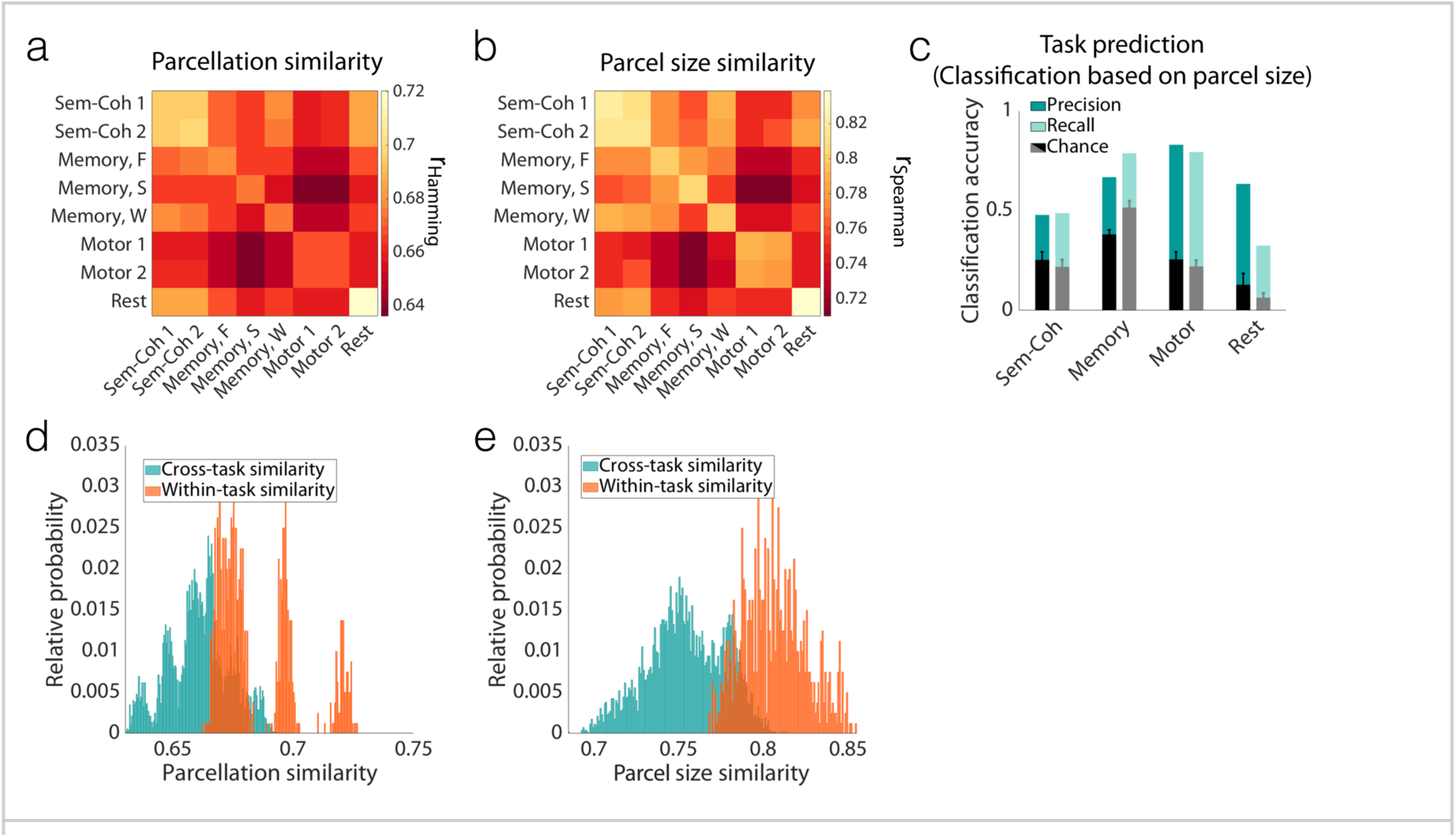
Replication of the finding that parcel definitions change with task condition; Midnight Scan Club (MSC) data. a) Pairwise parcellation similarity was calculated within and across functional conditions, averaged over all individuals. For every individual, voting-based ensemble analysis was used with 100 iterations. The matrix represents the average over all iterations. Similarity was assessed by r_Hamming_ = 1 – normalized Hamming distance. b) The same analysis as (a) was performed, this time using rank correlation of parcel-size vectors (r_Spearman_) as a proxy for parcellation similarities. c) The bars report the accuracy of predicting the functional condition using a leave-one-out cross-validated GBM classifier with parcel sizes as features. The predictive power is measured by the precision (dark cyan) and recall (light cyan) values for each condition. The precision (black) and recall (gray) values of 1000 null models are also reported (error bars represent ±s.d.). d) A histogram of parcellation similarities for all 100 iterations (averaged over all individuals) is depicted for within-condition (diagonal elements in [a]) and cross-condition (off-diagonal elements in [a]) comparisons. Sem-Coh 1 and Sem-Coh 2 are grouped into one condition, as are Motor 1 and Motor 2 conditions. The within- and cross-distributions are significantly different (K-S test; p<0.001). e) A histogram of parcel size similarities for all 100 iterations (averaged over individuals) is depicted for within-condition (diagonal elements in [b]) and cross-condition (off-diagonal elements in [b]) comparisons. The two distributions are significantly different (K-S test; p<0.001). Sem-Coh, semantic-coherence task; Memory F, S, and W, incidental memory task with faces, scenes, and words stimuli, respectively.

The results from the previous two data sets relied upon the construction of individual atlases across both sessions and conditions and were evaluated in terms of how the atlas *within an individual* changed between conditions. Next, we used the Human Connectome Project (HCP) 900 Subjects release (S900) data^33^ to determine if such condition-dependent reconfigurations could be observed when measured across multiple subjects introducing additional variance through inter-subject anatomic differences. The HCP data set includes task-based and resting-state fMRI data from 514 individuals (284 females; age = 22 – 36+), each of whom was scanned over a period of two days. Each scan session included seven tasks (day 1: working memory (WM), gambling, and motor tasks; day 2: language, social, relational, and emotion tasks) and 2 resting-state runs (one on each day).

We repeated the voting-based ensemble analysis described above, this time replacing sessions with subjects, such that we considered multiple subjects in multiple conditions and calculated the similarity between each pair of parcellations. Figure 5 demonstrates the pairwise parcellation similarities across subjects. Despite the introduction of inter-individual anatomic and functional organization variance, the primary finding that the parcellation map changes with condition remains highly significant (Figure 5a,d: fine-scale similarity [K-S test, p<0.001]; Figure 5b,e: coarse-scale similarity [K-S test, p<0.001]). As an even stronger assessment of generalizability, we repeated our previous analysis to predict, based on parcel size, the task during which the data were collected for previously unseen subjects. Given that we had a larger sample size (n=514), we further challenged our model by employing a more rigorous 10-fold cross-validated pipeline (rather than leave-one-out). We observed that our model could significantly predict, in novel subjects, the task administered while the data were acquired (Figure 5c, mean accuracy = 73%) based on parcel size, alone. This observation is an even stronger replication of our main findings; that is, the observed state-evoked parcellation reconfigurations are robust and reliable not only across different sessions, but also across distinct individuals.

**Figure 5.**
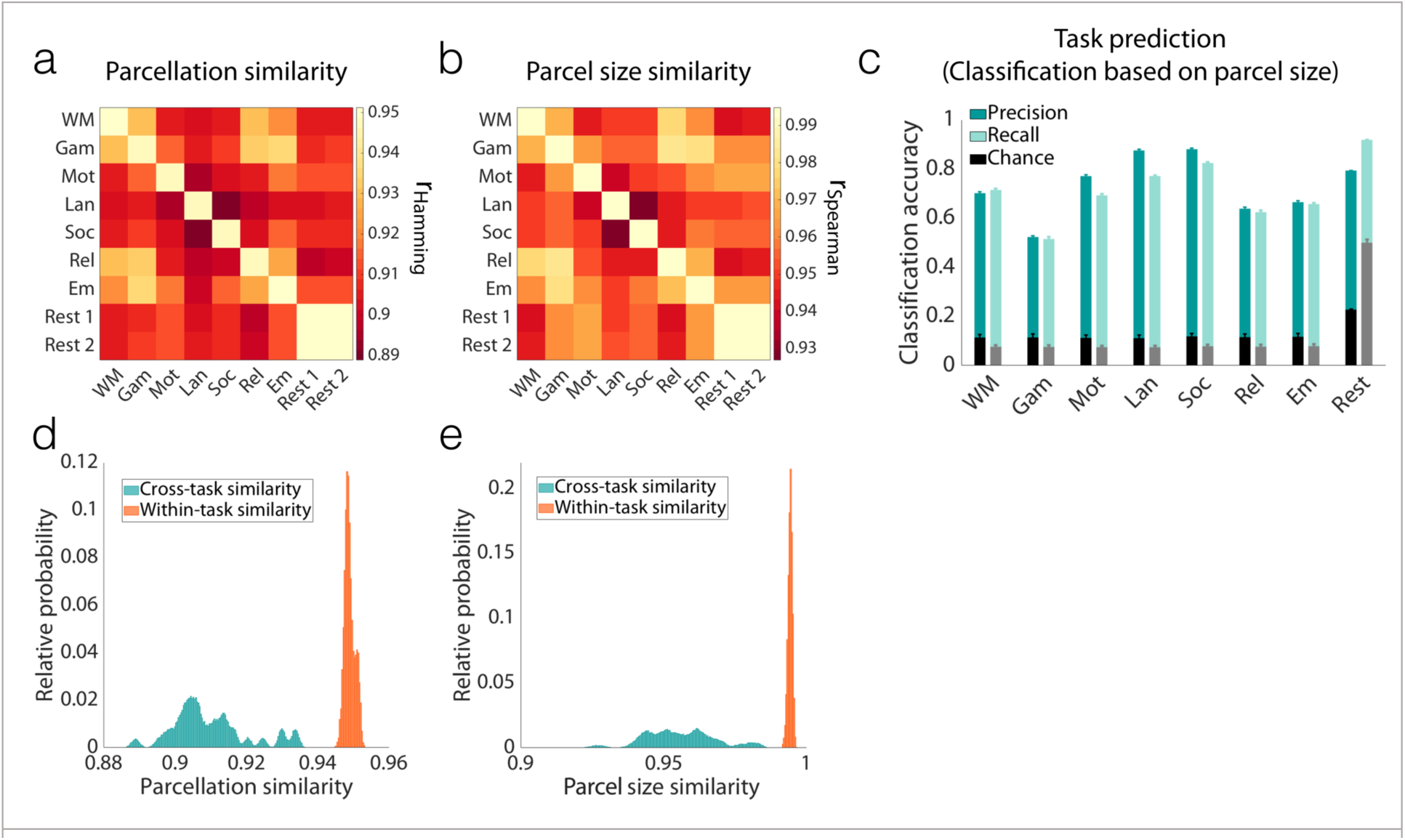
Replication of the finding that parcel definitions change with task condition, even when considered across individuals; Human Connectome Project (HCP) data. a) Pairwise parcellation similarity was calculated within and across functional conditions (9 conditions, N=514 subjects), using voting-based ensemble analysis with 1000 iterations. The matrix represents the average over all iterations. Similarity was assessed by r_Hamming_ = 1 – normalized Hamming distance. b) The same analysis as (a) was performed, this time using rank correlation of parcel-size vectors (r_Spearman_) as a proxy for parcellation similarities. c) The bars report the accuracy of predicting the functional condition using 10-fold cross-validated GBM classifier with parcel sizes as features, iterated 100 times. The predictive power is measured by precision (dark cyan) and recall (light cyan) values for each condition (reported as mean and s.d. across iterations). The precision (black) and recall (gray) values of the 1000 null models are also reported (error bars represent ±s.d.). d) A histogram of the parcellation similarities for all 1000 iterations is depicted for within-condition (diagonal elements in [a]) and cross-condition (off-diagonal elements in [a]) comparisons. Rest 1 and Rest 2 are grouped into one condition. The two distributions are significantly different (K-S test; p<0.001). e) A histogram of the parcel size similarities for all 1000 iterations is depicted for within-condition (diagonal elements in [b]) and cross-condition (off-diagonal elements in [b]) comparisons. The two distributions are significantly different (K-S test; p<0.001). WM, working-memory task; Gam, gambling task, Mot, motor task; Lan, language task; Soc, social task; Rel, relational task; Em, emotion task.

### Robustness of the state-evoked atlas reconfiguration across scales

In our final analysis, we sought to address the question of parcellation scale, asking if these reconfigurations are simply due to the rather large parcel sizes typically targeted in current parcellation approaches. If the parcel resolution were too low, then it could be the case that our parcels are composed of smaller subunits that do not change their shape but potentially exhibit altered connectivity with the other subunits dependent upon the functional state (hence modifying how they are grouped into a single parcel). If this is the case, then making atlases with more parcels should get to a level where the parcels remain unchanged even with changes in brain state. In the limit of parcels reduced to the size of a voxel, there can be no condition-induced change in parcel definition since the parcel is defined by the voxel size. Here we attempted to determine the critical resolution at which parcellations stabilize. To address this question, we repeated our analyses with Yale data, using atlases containing greater numbers of parcels, from 368, to 1041, to 5102 parcels. In each case, parcel reconfigurations were observed, even when the number of parcels was increased to 5102, more than ten times the typical number of parcels used in functional connectivity analysis^36-38^ (Figure 6). Results with 368-parcel and 1041-parcel atlases are reported in Figure S2 and Figure S3, respectively. To date, the human neuroscience community has not considered atlases with 5000 or more parcels.

**Figure 6.**
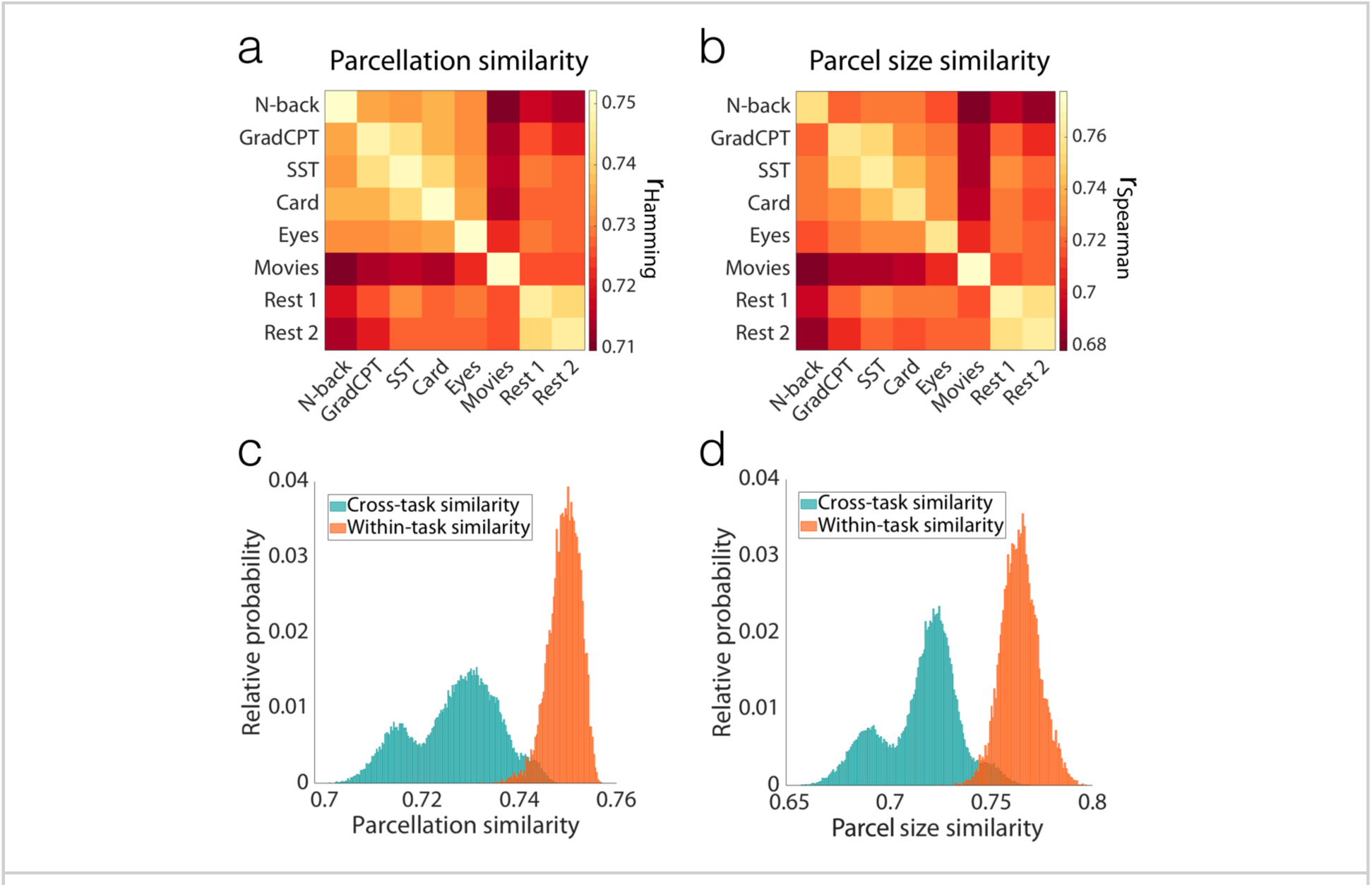
Parcel size effects: Even for an atlas with 5102 parcels (on average 25 voxels per parcel), parcel definitions change with task condition, with high reliability within conditions. a) Pairwise parcellation similarity was calculated within and across functional conditions, using voting-based ensemble analysis with 1000 iterations. Similarity was assessed by r_Hamming_ = 1 – normalized Hamming distance. b) The same analysis as (a) was performed, this time using rank correlation of parcel-size vectors (r_Spearman_) as a proxy for parcellation similarities. c) The histogram of the parcellation similarities for all 1000 iterations is depicted for within-condition (diagonal elements in [a]) and cross-condition (off-diagonal elements in [a]) comparisons. The two distributions are significantly different (K-S test; p<0.001). d) The histogram of the parcel size similarities for all 1000 iterations is depicted for within-condition (diagonal elements in [b]) and cross-condition (off-diagonal elements in [b]) comparisons. The two distributions are significantly different (K-S test; p<0.001).

## Discussion

Together, these findings suggest that there is no single functional atlas for the human brain. The boundaries of functionally defined parcels change with changes in brain state in a consistent and reproducible manner. These reconfigurations appear to be cognitively meaningful, as evidenced by their utility in predictive models of task condition and within-condition task performance.

### Reconfiguration of the connectome

These results are consistent with the extensive evidence for reconfigurations of the functional connectome. There is growing evidence that patterns of functional connectivity change with changing brain states (e.g., as induced by distinct tasks), and that these changes are functionally significant^39^. Recent work has highlighted individual differences in parcel spatial configuration and their impact on brain-behavior relationships^31^, but the common assumption that parcel boundaries are fixed for a given individual remains unchallenged. In fact, we show that while boundary reconfigurations may be relatively modest overall, they are highly task-specific, consistent, and functionally significant both within and across individuals, suggesting that tasks induce reliable perturbations of a core functional architecture of the human brain. Taking such reconfigurations into account may further inform efforts to relate changing patterns of functional connectivity to behavior, clinical symptoms, and cognition.

The stability of the state-evoked boundary reconfigurations across a large cohort of individuals, as demonstrated by the HCP data (Figure 5), is important. Previous work has shown that patterns of functional connectivity are unique to each individual^40^, and any state-evoked reconfigurations are also individual-specific^41^. That the average of these state-evoked reconfigurations across individuals does not mask out the effect of brain state is a significant finding, suggesting that task-induced parcel reconfigurations are not idiosyncratic and subject-specific, but rather robust and generalizable; that is, despite significant individual differences in brain functional organization, task-induced changes in this organization, as reflected in parcel boundary shifts, are highly similar across individuals.

### Composition of functional subunits

The cause of such parcel reconfigurations remains an important open question. At the atlas sizes considered here, any individual functional parcel will contain hundreds of thousands of neurons and may span multiple cortical subareas^42^, and a growing literature suggests that the BOLD signal can be used to identify inter-digitated neural representations, even within a circumscribed area with well-defined functional specialization^43-45^. And while it is possible that focal task activations affect parcel boundaries, they do not fully explain the present results, as parcel reconfiguration is broadly distributed across the brain, and change in parcel size relative to rest is not related to parcel task activation (Figure S6). Furthermore, the same results were obtained even after exclusion of voxels with significant task activations (Figure S7). Together, this suggests a more complex picture of parcels in flux across the brain, wherein different groups of neurons synchronize to execute different tasks resulting in different parcel definitions. It is possible that invariant boundaries may not be found until the acquisition resolution is of the order of a fundamental cortical unit, such as microcolumns (300 – 500 μm)^46,47^ or less. This presents an exciting opportunity for future work with high-field fMRI to interrogate the underpinnings of boundary shifts at high resolution.

The existence and nature of such a fundamental unit—and its potential impact on the BOLD signal—are open questions, but regardless of their answer, the findings here of substantial parcel reconfiguration even at atlas sizes of 5000 parcels (consisting of, on average, 25, 2 mm^3^ voxels, or ∼5.8 mm^3^ of tissue), suggest that current human functional parcellation atlases are far from this limit. Nevertheless, our goal does not have to be identifying parcels that correspond to a minimal functional or computational unit. In practice, a parcel that has a homogeneous, distinct identity in a given state is sufficient for many analyses. In other words, as long as the atlas is consistent for the state of interest, atlases with 200 – 400 parcels are entirely appropriate.

### Additional considerations

The exemplar-based parcellation approach used in this work imposes a constraint that holds the total number of parcels fixed and eliminates the problem of establishing correspondence between parcels under different conditions. Maintaining correspondence across parcellations facilitates quantitative assessment of the change in each parcel. It is possible that parcels not only reconfigure, but also blend and/or split, leading to a change in the total number of parcels with a change in brain state. This additional degree of freedom would make quantification of parcel changes across atlases obtained under different conditions more difficult due to the correspondence problem, while also potentially amplifying our main finding that there is no fixed functional atlas at the scale typically used in fMRI studies. In other words, without an exemplar-based approach, the reconfiguration of these parcels would likely be more extensive than that shown here.

The algorithm presented also constrains parcels to be contiguous. Without this constraint, it is possible for connectivity-based parcellations to group voxels, that may be widely separated spatially, into single parcels with very high homogeneity in their time-courses. If parcels are allowed to be non-contiguous, it becomes an open question as to how much distance is allowed between the different parts of a parcel. With no consideration given to spatial contiguity, a successful parcellation algorithm could simply rank voxel time-courses by similarity and then split this ranking into *n* parcels, which would provide maximal parcel homogeneity (a common measure of parcellation success), but would identify broadly distributed brain networks rather than functionally specialized, circumscribed brain regions.

The present work focuses on defining parcels with hard boundaries. However, one can use the voxel uncertainties to define soft borders, as displayed in the blue boundaries in Figure 3. One can also define a probabilistic soft parcellation by considering the consistency of voxel-to-parcel assignments across different functional conditions. While defining such soft borders is feasible, most parcellation approaches to date have focused on hard boundaries. One application of such functional atlases is in building connectivity matrices, and with soft boundaries the question of how to weight the time-courses across these boundaries is currently unanswered. Given the use of parcellations with hard boundaries for validated, commonly used functional connectivity analysis pipelines, we chose to focus here on such hard boundaries.

States are defined in this work as task conditions with acquisitions spanning a series of continuous performance, event-related, and blocked tasks. It is likely that parcel reconfiguration occurs over considerably shorter periods of time than these minutes-long task intervals, particularly given the growing literature on the dynamic nature of functional brain organization^48^ and the present finding that there are meaningful, state-induced changes in parcellation boundaries within a given task (i.e., that task performance can be predicted from parcel size). Future work may seek to characterize how parcellation boundaries shift to reflect the dynamic reconfiguration of macroscale neural circuitry underlying moment-to-moment changes in brain state.

This work also does not invalidate multimodal parcellation approaches that combine both anatomical and functional data (including functional data across a wide range of task and resting states), but suggests that such approaches, by defining a mean atlas across states, may mask meaningful and informative functional reconfigurations associated with specific brain states.

### Implications

Essentially all publications to date that employ parcellations with hard boundaries have used fixed atlases at the group or individual level, with parcel boundaries defined anatomically or functionally (or through a combination of methods). Such work assumes that parcel boundaries do not change meaningfully for a given individual or group, but the findings here suggest that this is not the case for atlases defined using functional MRI data. While functional parcel reconfigurations would not affect results generated using data collected during a single state, they may affect investigations of connectivity changes across states. More generally, it suggests that a single functional parcellation applicable to all brain states is not appropriate. Further, these findings suggest that state-dependent changes in connectivity could be attributable in part to reconfiguration of the underlying parcellation, and this should be considered as part of the interpretation of changes in connectivity that occur across different cognitive conditions.

It has been posited that imposing an atlas on the brain is simply a data-reduction strategy to reduce the connectivity matrix to a manageable size. This may indeed be a useful feature (for a discussion, see Eickhoff et al.^20^), and this work does not impact its utility. The findings here, however, support the notion that function-based parcellations are not only a means of data dimensionality reduction but also a way to reveal meaningful patterns of, and changes in, brain functional organization. The results demonstrating prediction of state and performance based on the particular parcel size support this conclusion.

New approaches to individualized parcellation, such as the approach used in this work, could potentially lead to custom state-dependent atlases for individuals or groups, which may in turn provide further insight into understanding state-dependent functional reorganization of the human brain. Much work needs to be done to understand the relationship between functional edge strength measures and these variable parcel configurations, and the impact of these variables on functional connectivity measures in both health and disease. It is an open question whether fixed functional subunits, invariant to brain state, can be defined in the human cortex with current state-of-the-art neuroimaging methods. We do know from these results however, that such an atlas would need to have more than 5000 parcels. Nevertheless, state-dependent changes in parcellation boundaries offer an important tool to study neural representation and dynamic interactions in the human brain.

## Conclusion

This work demonstrates that there is no single functional atlas for the human brain at the 200 – 5000 parcel resolution level, but rather that parcels reconfigure depending on task-induced state in a robust and reliable manner. Such reconfigurations are distinct and reliable enough to use quantitative parcellation characteristics (i.e., parcel size) to predict task condition across multiple tasks, as well as within-task performance. That functional parcel definitions are fluid must be considered when interpreting changes in functional connectivity patterns across states. Such parcel reconfigurations may be leveraged to better understand dynamic changes in functional organization of the human brain. These results therefore provide another mechanism by which to understand the flexible brain and how it reconfigures to perform the task at hand. The derivation of state-specific, individualized functional atlases could provide an important tool for human neuroscience, and this work calls for the continued development and validation of such approaches.

## Materials and Methods

Three data sets were used in this work. The primary data were collected at Yale University from one healthy subject (author R.T.C.). The validation data were obtained from the publicly available Midnight Scan Club (MSC) data set^49^, and Human Connectome Project (HCP). These three data sets are described in detail below.

### Yale Data

#### Participant and processing

The primary subject R.T.C. is a healthy left-handed male, aged 56 years old at the onset of the study. The subject provided written informed consent in accordance with a protocol approved by the Human Research Protection Program of Yale University.

The subject was scanned at Yale University 33 times (that is, 33 sessions) over ten months. Scans were typically performed on Wednesdays at 8:30 am and Fridays at 2:00 pm. Functional MRI data were acquired on 2 identically configured Siemens 3T Prisma scanners equipped with a 64-channel head coil at the Yale Magnetic Resonance Research Center. Three sessions were excluded from analysis: the first two sessions were excluded because of the considerable adjustment in task design after session 2, and the ninth session was excluded due to interruptions in task presentation due to interruptions in task presentation.

The first session was used to acquire structural MRI data. High-resolution T1-weighted 3D anatomical scans were performed using a magnetization prepared rapid gradient echo (MPRAGE) sequence with the following parameters: 208 contiguous slices acquired in the sagittal plane, repetition time (TR) = 2400 ms, echo time (TE) = 1.22 ms, flip angle = 8°, slice thickness = 1 mm, in-plane resolution = 1 mm × 1 mm, matrix size = 256 × 256. A T1-weighted 2D anatomical scan was acquired using a fast low angle shot (FLASH) sequence with the following parameters: 75 contiguous slices acquired in the axial-oblique plane parallel to AC-PC line, TR = 440 ms, TE = 2.61 ms, flip angle = 70°, slice thickness = 2 mm, in-plane resolution = 0.9 mm × 0.9 mm, matrix size = 256 × 256.

Functional scans were performed using a multiband gradient echo-planar imaging (EPI) pulse sequence with the following parameters: 75 contiguous slices acquired in the axial-oblique plane parallel to AC-PC line, TR = 1000 ms, TE = 30 ms, flip angle = 55°, slice thickness = 2 mm, multiband acceleration factor = 5, in-plane resolution = 2 mm × 2 mm, matrix size = 110 × 110.

Data were analyzed using BioImage Suite^50^ and custom scripts in MATLAB (MathWorks). Motion correction was performed using SPM (https://www.fil.ion.ucl.ac.uk/spm/). White matter and CSF masks were defined in MNI space and warped to the single-subject space using a series of linear and non-linear transformations (see Scheinost et al.^51^). The following noise covariates were regressed from the data: linear, quadratic, and cubic drift, a 24-parameter model of motion^52^, mean cerebrospinal fluid signal, mean white matter signal, and mean global signal. Finally, data were temporally smoothed with a Gaussian filter (*σ* = 1.55).

#### Dimensional task battery design

Functional scans were 6 minutes 49 seconds each, including initial shim and 8s discarded acquisitions before the start of each task. Tasks varied slightly in length (see below), but were all approximately 6 minutes in duration. A fixation cross was displayed after the end of each task and lasted until the beginning of the next task. Each task, with the exception of movie watching, was preceded by instructions and practice, after which the subject had the opportunity to ask questions before the scan began. All responses were recorded using a 2×2 button box.

Each session consisted of two resting-state runs and six task runs. The first and last functional runs (runs 1 and 8) were resting-state runs, during which the participant was instructed to stay still with his eyes open. Runs 2 – 7 were task runs, with the order counterbalanced across sessions.

### Task details

#### N-Back Task

The *n*-back task was adapted from that used in Rosenberg et al.^24^. In this task, the participant was presented with a sequence of images and was instructed to respond via button press if the image was different than the image presented two before, and to withhold response if it was the same. Images were presented for 1 second, followed by a 1-second inter-trial interval (ITI; fixation cross). The target (i.e., matching image) probability was 10%. There were two blocks, each with 90 trials. One block used images of emotional faces and the other block used images of scenes^53,54^. Block and stimulus order were randomized for each session. Task performance was assessed by *sensitivity* (*d’*), defined as hit rate relative to false alarm rate^26^.

#### Gradual-onset Continuous Performance Task (gradCPT)

The gradCPT task was adapted from that described in Esterman et al.^25^ and Rosenberg et al.^26,27^. In this task, the participant viewed a sequence of 450 scenes (city or mountain) that gradually transitioned via linear pixel-by-pixel interpolation from one to the next over 800ms. The participant was instructed to respond via button press to cities and to withhold response to mountains. Stimulus order was randomized, and 10% of images were mountains. Task performance was assessed by *sensitivity* (*d*′).

#### Stop-Signal Task (SST)

The stop-signal task was adapted from that implemented in Verbruggen et al.^28^. In this task, the participant was required to determine via button press whether a presented arrow was pointing left (right index finger) or right (right middle finger). On 25% of trials, the arrow turned blue after some delay, indicating that the participant should withhold response. This stop-signal delay (SSD) was initially set to 250ms, and was continuously adjusted via the staircase tracking procedure (50ms increase after correct inhibition trials; 50ms decrease after each failure to inhibit). The arrow was presented for 1.5 seconds, followed by 0.5 seconds of fixation; there were 176 trials in total, with stimulus order randomized within block. Task performance was assessed by *missing probability*, defined as the percentage of missed responses on no-signal trials^28^.

#### Card Guessing Task

The card guessing task was adapted from that originally developed by Delgado et al.^29^ and subsequently extended^55,56^. In this task, the participant was presented with a card and asked to guess if the number on the back was lower than 5, or greater than 5 but less than 10. The question mark card was displayed for 1.5 seconds, or until the participant responded (right index finger for “lower,” right middle finger for “higher”). The card then “flipped over” to reveal the number. The number was displayed for 0.5 seconds, followed by an arrow for 0.5 seconds to indicate accuracy (green and up for correct, red and down for incorrect), which was in turn followed by a 1-second inter-trial interval (fixation cross). There were 10 blocks, each with 10 trials, and guess accuracy was deterministic, such that in half of the blocks (“high win”), the participant was correct 70% of the time, while in the other half of the blocks (“high loss”) he was correct 30% of the time; block (high win/loss) and trial (correct/incorrect) orders were randomized. Task performance was assessed by *RT variability*, defined as standard deviation of reaction time^57^.

#### Reading the Mind in the Eyes Task (“Eyes Task”)

The Eyes Task was adapted from that originally described in Baron-Cohen et al.^30^. In this task, the participant viewed a series of photographs of an individual’s eyes with four “mental state terms”^30^, one in each corner of the image, and was instructed to select via button press (with each button corresponding to one corner, and thus one term) the term that best described what the individual was thinking or feeling. There were 36 images in total. Each was presented once, in random order, for 9.25 seconds or until the participant responded; the remainder of each 10-second trial consisted of a fixation cross. Task performance was assessed by *RT variability*, defined as standard deviation of reaction time for correct trials.

#### Movies Task

In this task, three movie clips were presented in continuous series; each was approximately 2 minutes long. The first clip was a trailer for “Inside Out,” the second clip was the wedding scene from “Princess Bride,” and the third clip was a trailer for “Up;” order was fixed across sessions. The participant was instructed to relax and enjoy the movies; no responses were required. No task performance was recorded.

### Midnight Scan Club (MSC) Data

#### Participants and processing

The MSC data set^49^ includes data from 10 healthy individuals (5 females; age = 24 – 34); each underwent 1.5 hours of functional MRI scanning on 10 consecutive days, beginning at midnight. For details of the data acquisition parameters and sample demographics see Gordon et al^49^. Two individuals were excluded from the analysis: MSC08 was excluded because of excessive head motion and self-reported sleep^49^; MSC10 was excluded for insufficient data (missing one session of incidental memory task).

All data were preprocessed using BioImage Suite^50^. Data were transformed to MNI space to facilitate analysis across multiple subjects. Preprocessing steps included regressing 24 motion parameters, regressing the mean time courses of the white matter and cerebrospinal fluid as well as the global signal, removing the linear trend, and low pass filtering.

#### Task battery design

Each scanning session started with a 30-min resting-state fMRI scan, followed by three separate task-based fMRI scans: motor task (2 runs per session, 7.8 min combined), incidental memory task (3 runs per session, 13.1 min combined), and semantic-coherence task (2 runs per session, 14.2 min combined).

#### Motor Task

The motor task was adapted from that used in the Human Connectome Project (HCP)^55^. In this task, participants were cued to perform one of the following movements: closing/relaxing their hands, flexing/relaxing their toes, or wiggling their tongue. Each block started with a 2.2 s cue indicating which movement to perform, followed by a central caret (flickering every 1.1s) to signal the movement. Each run consisted of 2 blocks of each movement type and 3 blocks of resting-fixation (15.4 s total).

#### Incidental Memory Task

The incidental memory task consisted of three different types of stimuli (scenes, faces, and words), each presented in a separate run. For scene runs, participants were asked to decide if the presented scene was indoors or outdoors. For face runs, participant made male/female judgments. For word runs, participants made abstract/concrete judgments. Each run consisted of 24 stimuli, each repeating 3 times. Stimuli were presented for 1.7 s with a jittered 0.5-4.9 s inter-stimulus interval. All stimuli were taken from publicly available sources (see Gordon et al.^49^ for details).

#### Semantic-Coherence Task

The semantic-coherence task had a mixed block/event-related design, consisting of two different conditions (“semantic” and “coherence”). In the “coherence” task, participants viewed a concentric dot pattern (Glass^58^) with 0% or 50% coherence, and made binary decisions whether the pattern was concentric or random. In the semantic task, participants viewed a word and indicated whether the word is a noun or verb. Each run consisted of two blocks of each task, separated by 44 s of rest. Each block started with a 2.2 s cue indicating which task was to be performed. Blocks consisted of 30 trials. Stimuli were presented for 0.5 s with a variable 1.7-8.3 s ISI. Each block finished with a 2.2 s cue indicating the end of the task block.

### Human Connectome Project (HCP) Data

#### Participants and processing

The HCP data set^59^ includes data from 897 healthy individuals (S900) scanned during nine functional conditions (seven tasks and two rest). For details of the data acquisition parameters see Ugurbil et al.^60^ and Smith et al^61^. Analyses were restricted to subjects for whom data were available for all nine functional conditions (with left-right (LR) and right-left (RL) phase encoding). To mitigate the substantial effects of head motion on functional parcellations, we further excluded subjects with excessive head motion (defined as mean frame-to-frame displacement > 0.1 mm and maximum frame-to-frame displacement > 0.15 mm), leaving 514 subjects (284 females; age = 22 – 36+) for analysis.

The HCP minimal preprocessing pipeline was employed^62^, which includes artifact removal, motion correction and registration to MNI space. Further preprocessing steps were performed using BioImage Suite^50^ and included standard preprocessing procedures^40^ including regressing 24 motion parameters, regressing the mean time courses of the white matter and cerebrospinal fluid as well as the global signal, removing the linear trend, and low pass filtering.

#### Task battery design

Functional MRI scans were acquired during two different days: Day 1 included two runs (LR and RL) of the working memory (WM) task (5:01 min per run), incentive processing (gambling) task (3:12 min), motor task (3:34 min), and rest (14:33 min); day 2 included two runs of the language processing task (3:57 min), social cognition (theory of mind) task (3:27 min), relational processing task (2:56 min), emotion processing task (2:16 min), and rest (14:33 min).

The details of task design have been previously described^55,59^. We provide a brief description of each task and an overview of the relevant aspects below.

#### Working Memory Task

In this task, participants performed a visual *n*-back task, with blocked 0-back and 2-back conditions using four stimulus categories (faces, places, tools, body parts). Each run consisted of 8 task blocks (10 trials each), with each stimulus category used twice, and 4 fixation blocks. Each block started with a 2.5 s cue indicating the task type (0-back versus 2-back) and the target (for 0-back).

#### Gambling Task

In this task, participants were presented with a mystery card and asked to guess if the number on the back was lower than 5, or greater than 5 but less than 10. On reward trials, participants were shown the number on the card, a green up arrow, and “$1”; on loss trials, participants were shown the number on the card, a red down arrow, and “-$0.50”; on neutral trials, participants were shown the number 5 and a gray, double-headed arrow. Each run consisted of 4 task blocks (8 trials each) and 4 fixation blocks. In half of the blocks (“mostly reward”), subjects were correct in 6 out of 8 trials (the remaining 2 trials were either neutral or loss), while in the other half of the blocks (“mostly loss”) they were incorrect in 6 out of 8 trials (the remaining 2 trials were either neutral or reward).

#### Motor Task

Participants were presented with visual cues that asked them to tap their left or right fingers, squeeze their left or right toes, or move their tongue. Each block started with a 3 s cue indicating which movement to perform. Each run consisted of 2 blocks of tongue movements, 4 blocks of hand movements (2 left and 2 right), 4 blocks of foot movements (2 left and 2 right), and 3 blocks of resting-fixation.

#### Language Task

In this task, participants were aurally presented with 4 blocks of a story task and 4 blocks of a math task. In the story task, they heard brief fables (5-9 sentences) and completed two-alternative forced-choice questions about the topic of the story. In the math task, they completed addition and subtraction problems in a two-alternative forced-choice setting.

#### Social Task

In this task, participants were presented with 20-s video clips of objects (squares, circles, triangles) either interacting (theory-of-mind) or moving randomly. Participants were asked to choose between three potential responses (“mental interaction”, “no mental interaction”, and “not sure”). Each run consisted of 5 video blocks (2 mental and 3 random in one run, 3 mental and 2 random in the other run) and 5 fixation blocks.

#### Relational Task

The relational task consisted of two different conditions (“relational” and “matching”). In the relational condition, participants were presented with two pairs of objects with one pair at the top and the other pair at the bottom of the screen. They were asked to decide whether the bottom pair of objects differed along the same dimension (i.e., shape or texture) as the top pair. In the control matching condition, they were presented with two objects at the top, one object at the bottom, and a word (“shape” or “texture”) in the middle of the screen. They were asked to determine whether the bottom object matched either of the top two objects on the dimension specified by the word. Each run consisted of 3 relational blocks (4 trials each), 3 matching blocks (5 trials each) and 3 fixation blocks.

#### Emotion Task

In this task participants were presented with blocks of “face” and “shape” tasks, and were asked to determine which of the two faces (or shapes) presented at the bottom of the screen matched the face (or shape) at the top of the screen. Faces had either angry or fearful expressions. Each block started with a 3 s cue indicating which task to perform. Each run included 3 face blocks (6 trials each) and 3 shape blocks (6 trials each), with 8 s fixation at the end of each run.

#### Individualized and state-specific parcellation algorithm

Here, we extended our previously developed individualized parcellation algorithm^11^ to account for the spatial contiguity of the parcels at the individual level. The presented algorithm is a priority-based submodular method that defines functional parcels in a streaming fashion, for every individual in every functional state.

Our algorithm runs in three steps:

i. **Registration of the initial group-level parcellation.** In the first step, an off-the-shelf group-level parcellation is applied to each individual’s data, assigning each voxel to a parcel defined by the group parcellation. At this step, all individuals in every state have the same parcel definitions.
ii. **Exemplar identification.** In the second step, for every group-defined parcel in an individual brain, an exemplar is identified by maximizing a monotone nonnegative submodular function (see Eq. 2).
iii. **Spatially-constrained voxel-to-parcel assignment.** Finally, the third step assigns every voxel in each individual brain to the functionally closest exemplar while ensuring the spatial contiguity of the resulting parcel.

A visual illustration of the parcellation algorithm is provided in Figure 7a.

#### Exemplar identification

The exemplar identification algorithm can be viewed as a data summarization step, where the goal is to summarize a massive amount of data by fewer representative points or exemplars. A classic way of defining such exemplars is by finding the set *S* that minimizes the following loss function, subject to the constraint |*S*| = *k* (known as *k*-medoid problem).

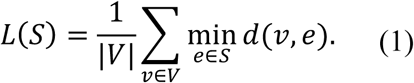

In this equation, *V* is the ground set consisting of all data points, *d*: *V* × *V* → *R* is a dissimilarity function defined on every pair of data points, and *S* is the objective exemplar set. Intuitively, *L*(*S*) measures how much information we lose if we summarize the entire ground set to the exemplar set by representing each data point with its closest exemplar.

Minimizing this loss function (1) is NP-hard, as it requires exponentially many inquiries. Using an appropriate auxiliary exemplar *ν*_0_, we transform the minimization of (1) into the maximization of a non-negative monotone submodular function^63^, for which general greedy algorithms provide an efficient 1 - 1/e ≈ 0.63 approximation to the optimal solution^64^:

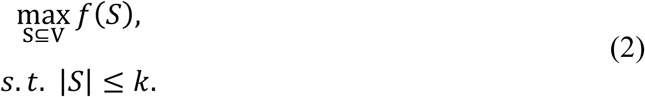

Where:

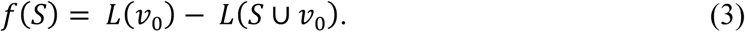

In practice, the greedy algorithm provides a considerably closer approximation to the optimal solution (see Salehi et al.^34^). For the choice of auxiliary exemplar, any vector *ν*_0_ whose distance to every data point is greater than the pairwise distances between data points can be used.

##### Definition 1 (Submodularity)

A function *f*: 2^*V*^ → *R* is submodular if for every *A* ⊆ *B* ⊆ *V* and *e* ∈ *V*\*B* it holds that *f*(*A* ∪ *e*) – *f*(*A*) ≥ *f*(*B* ∪ *e*) – *f*(*B*). That is, adding an element *e* to a set *A* increases the utility at more than (or at least equal to) adding it to *A*’s superset, *B*, suggesting natural diminishing returns.

#### Spatially-constrained voxel-to-parcel assignment

After identification of all exemplars (one per parcel), every voxel in the individual brain is assigned to the functionally closest exemplar while taking the spatial contiguity of the parcel into account. The spatial contiguity is assured by utilizing priority queues. Every exemplar *i* is assigned a priority queue (denoted as *q*_*i*_). Initially, all the queues are empty. In the first round, spatial neighbors of exemplar *i* are pushed into *q*_*i*_. The voxels in each queue are sorted according to their functional distance to the corresponding exemplar such that the voxel with minimum functional distance (maximum similarity) is in front. Next, the front voxel in each *q*_*i*_ is considered as a potential candidate for being assigned the label *i*. Among all these candidates, the one with minimum distance to its corresponding exemplar is selected and assigned the exemplar’s label. Next, this voxel is popped out of the queue, and all of its spatial neighbors are pushed into the same queue. The algorithm continues until all the voxels are assigned a label. Note that at every step of the algorithm the labeled voxel is ensured to be spatially connected to its exemplar (either directly or through other previously labeled voxels).

The quality of the proposed parcellation approach was assessed and compared with the initial group-level parcellation, using two internal clustering validation methods: Homogeneity (Figure 7b, top panel) and Davies-Bouldin index (DB) (Figure 7b, bottom panel).

#### Implementation details

Here, we implemented an accelerated version of the greedy algorithm, called lazy greedy^65^. Also, here *k* = 1, as we attempt to identify one exemplar per parcel (see Salehi et al.^11^ for details of interpretation and alternative approaches). For the choice of dissimilarity measure, we used squared Euclidean distance, after normalizing all the voxel-level time courses to a unit norm sphere centered at the origin. A point with the norm greater than 2 was used as the auxiliary exemplar. The parcellation algorithm was applied to each fMRI run (each individual in each state) independently, and thus was efficiently employed through parallelization. For HCP data, we restricted our analysis to left-right (LR) phase encoding.

#### Initial group-level parcellations

As the initial group-level parcellation, we primarily used a 268-parcel atlas, previously defined in our lab using a spectral clustering algorithm on resting-state data of a healthy population^15,40^. We replicated the results with more fine-grained atlases including 368, 1041, and 5102 parcels. The 368-parcel atlas was defined by integrating the parcellation of cortex from Shen et al.^15^, subcortex from the anatomical Yale Brodmann Atlas^66^, and cerebellum from Yeo et al.^67^. Similarly, the 1041-parcel parcellation was defined by integrating the subcortical and cerebellum portion of the 368-parcel parcellation with the 1000-parcel cortex parcellation from Yeo et al^67^. To define the 5102-parcel atlas, we started from the 1041-parcel parcellation and randomly divided all the parcels until the number of voxels per parcel reached approximately 25.

#### Statistical voting-based ensemble analysis

To estimate the similarity of parcellations within and across states, we employed a voting-based ensemble analysis. The reason for this analysis is two-fold: First, to help rule out noise and session effects as potential confounds; and second, to aid with the statistical comparison across states by generating a large distribution of state-specific parcellations. For each condition, we divided all sessions (for Yale and MSC data) or individuals (for HCP data) into two equal-size groups: group 1 and group 2. We took the relative majority vote over the parcellations of each group by assigning each voxel to the parcel for which the maximum number of sessions voted. This resulted in two parcellations for each functional state, one for each group. Next, we assessed the similarity between every pair of the non-overlapping parcellations both within and across states. For instance, if there are *m* functional states, this analysis generates an *m* × *m* matrix, where each element (*i, j*) represents the similarity between the parcellation of group 1 for state *i* and the parcellation of group 2 for state *j*. Thus, the diagonal elements represent the within-state similarities while the off-diagonal elements represent the cross-state similarities. We repeated the entire analysis 1000 times, generating an ensemble of *m* × *m* similarity matrices. The normalized distribution of the within-state and cross-state similarity values were depicted as histograms, and compared using the non-parametric Kolmogorov–Smirnov test (Figures 2d, e, 4d,e, and 5d,e). The averages of these similarity matrices were also displayed (Figures 2a,b, 4a,b, and 5a,b).

#### Similarity measures between parcellations

We compared parcellations at two different scales: at the fine scale, we studied the ratio of voxels that change their parcel assignment across different parcellations. The fine-scale similarity was calculated using 1 – normalized Hamming distance (Figures 2a, 4a, 5a). At the coarse scale, we studied the changes in the parcel sizes across parcellations. The coarse-scale similarity was computed using Spearman correlation between parcel-size vectors (Figures 2b, 4b, 5b).

#### Functional state decoding using parcel size as feature

We established a cross-validated predictive model that predicts the functional state of each unseen sample solely based on the size of parcels in that parcellation. Using a *k*-fold cross-validated approach, we trained and tested a gradient boosting classifier (GBM; with 300 estimators and learning rate = 0.1) using parcel sizes as features and the functional state as output. We randomly divided the entire data into *k* folds (*k*=*n* for the Yale and MSC data sets, where *n* is the total number of runs, and *k*=10 for the HCP data set). At each step, the model was trained on *k*-1 folds and tested to predict the state of the left-out fold. The predictive power of the model was estimated using precision (also known as positive predictive value) and recall (also known as sensitivity), calculated separately for each state. Both precision and recall measures range between 0 and 1, with higher values indicating higher predictive power. Precision calculates what fraction of the retrieved instances were actually relevant, while recall expresses what fraction of relevant instances were retrieved. In the case of 10-fold cross-validation (i.e., for the HCP data set), we repeated the predictive analysis 100 times to account for the randomness of the folds, and reported the precision and recall measures as the mean and standard deviation across all iterations.

To evaluate the significance of the results, we employed non-parametric permutation testing: we randomly permuted the output vector (here the functional states) 1000 times, and each time ran the permuted values through the same predictive pipeline and calculated the precision and recall measures of the permuted states.

#### Task performance prediction using parcel size as feature

To establish the relevance of parcellation boundary changes to within-task variation in brain state, we developed a cross-validated predictive pipeline that predicts task performance during each Yale data session from the parcel sizes in the parcellation for that session. Using a leave-one-out cross-validated approach, we trained and tested a ridge regression model (with regularization parameter = 1), using parcel sizes as features and the performance scores as output. For each task, a model was trained on *n-*1 sessions (of that task) and used to predict performance for the left-out session. Analyses were performed for each task independently. To measure performance, we used *d’* for *n*-back and gradCPT tasks, *RT variability* for eyes and card-guessing tasks, and *missing probability* for SST. The predictive power was estimated using r-values, defined as the square root of the percentage of the explained variance. To estimate the significance of the results, we employed non-parametric permutation testing, where we randomly permuted the behavioral scores 1000 times, and each time ran the permuted scores through the predictive pipeline and calculated the r-value.

#### Effects of parcellation algorithm on the state-evoked parcel reconfigurations

To ensure that the observed state-evoked reconfigurations did not depend on the choice of parcellation, we employed two additional experiments. First, we repeated our analysis using a slight modification of our exemplar-based algorithm, where instead of employing a data-driven exemplar selection we manually fixed the exemplar to be a random voxel in the parcel, with a particular exemplar for a given parcel fixed across all sessions and conditions. This means the initial reference voxel was identical for a given parcel across all sessions and conditions. In this case, only the time-course data could shift the parcellation since all parcellations started from the same exemplar voxels across conditions and sessions. Results remained largely unchanged (Figure S4). Second, we replicated our results using Wang et al.’s individualized parcellation algorithm, which is based on an iterative *k*-means clustering approach^35^. Wang’s method has an averaging step for which we used a standard uniformly weighted averaging. For results, see Figure S5.

#### Effects of task activation on the state-evoked parcel reconfigurations

To address the question of whether the parcellation reconfigurations were driven by task activation, we performed two experiments. First, we tested whether the mean task activation (calculated per parcel) correlates with differences in parcel size relative to rest (Figure S6). Next, we eliminated parcels with significant task activation for any of the tasks and repeated the entire analysis (Figure S7). Significance was defined as |z|>1.96, associated with 95% confidence interval or p<0.05. These two experiments were performed on both Yale and HCP data sets. For Yale data, because all the tasks were continuous performance tasks, we approximated the task activations per parcel by computing the difference between the average temporal signal during task and rest. For HCP data, we generated task effect size maps using all available individuals’ volume-based, FEAT-analyzed, first-level GLM output (COPE files) from the 1200 Subjects Release (S1200) for a given task to generate, using FSL FEAT’s FLAME (FMRIB’s Local Analysis of Mixed Effects^68^), cross-subject, voxel-wise Cohen’s d effect-size contrast maps from t-statistic maps. We used the following contrasts for each task: REWARD-PUNISH (Gambling, cope6), 2BK-0BK (WM, cope11), FACES-SHAPES (Emotion, cope3), STORY-MATH (Language, cope4), AVG (Motor, cope7), REL-MATCH (Relational, cope4), and TOM-RANDOM (Social, cope6). We then applied the initial 268-parcel group-level parcellation^15^ to these voxel-level maps to calculate a mean task effect size per parcel for each of the tasks.

#### Effects of head motion on the state-evoked parcel reconfigurations

To rule out the possibility that state-evoked reconfigurations were simply driven by characteristic head motion patterns specific to each functional condition, we tested whether there is significant difference in head motion between different functional conditions. We performed pairwise Wilcoxon signed-rank test (which is a non-parametric statistical hypothesis test to compare two dependent samples) on the mean frame-to-frame displacement, and corrected for multiple comparison using Bonferroni correction (Figure S8).

## Code and Data Availability

All the functional parcellations are available online on the BioImage Suite NITRC page (https://www.nitrc.org/frs/?group_id=51). An interactive visualization of the state-specific parcellations can be found here: http://htmlpreview.github.io/? https://github.com/YaleMRRC/Node-Parcellation/blob/master/Parcellation_visualization.html. MATLAB, C++ (for parcellation algorithm), and Python (for predictive modeling) scripts were written to perform the analyses described; these codes are available on GitHub at https://github.com/YaleMRRC/Node-Parcellation.git. The force-directed graph visualization (R script) is released separately under the terms of GNU General Public License and can be found here: https://github.com/YaleMRRC/Network-Visualization.git. Three data sets were used to support the findings of this study. Yale data are publicly available at International Neuroimaging Data-sharing Initiative (INDI) [http://fcon_1000.projects.nitrc.org/]. Midnight Scan Club (MSC) data are publicly available through the Open fMRI data repository at https://openneuro.org/datasets/ds000224/versions/00002. Human Connectome Project (HCP) data (S900) are publicly available at https://www.humanconnectome.org/study/hcp-young-adult/document/900-subjects-data-release.

## Acknowledgments

This work was supported by NIH Grants R01 MH111424 (R.T.C.) and DARPA Young Faculty Award D16AP00046 (A.K.). Two publicly available data sets were used in this work. The first data set was provided by the Midnight Scan Club (MSC)^49^ project, funded by NIH Grants NS088590, TR000448 (NUFD), MH104592 (DJG), and HD087011 (to the Intellectual and Developmental Disabilities Research Center at Washington University); the Jacobs Foundation (NUFD); the Child Neurology Foundation (NUFD); the McDonnell Center for Systems Neuroscience (NUFD, BLS); the Mallinckrodt Institute of Radiology (NUFD); the Hope Center for Neurological Disorders (NUFD, BLS, SEP); and Dart Neuroscience LLC. This data was obtained from the OpenfMRI database. Its accession number is ds000224. The second data set was provided by the Human Connectome Project, WU-Minn Consortium (Principal Investigators: David Van Essen and Kamil Ugurbil; 1U54MH091657) funded by the 16 NIH Institutes and Centers that support the NIH Blueprint for Neuroscience Research; and by the McDonnell Center for Systems Neuroscience at Washington University.

## Author Contributions

M.S., A.K., D.S., and R.T.C. conceived and formulated the study. M.S. and A.K. developed the spatially-constrained exemplar-based submodular parcellation algorithm. M.S. performed the parcellation analyses, the statistical analyses, and the predictive modeling. A.S.G. designed the task battery, collected, and preprocessed the primary data (RTC data set). M.S. preprocessed MSC data set. M.S. wrote the manuscript with contributions from R.T.C. and A.S.G. All other authors commented on the paper.

## References

1 Eickhoff, S. B., Constable, R. T. & Yeo, B. T. Topographic organization of the cerebral cortex and brain cartography. NeuroImage, doi:10.1016/j.neuroimage.2017.02.018(2017).

2 Brodmann, K. Vergleichende Lokalisationslehre der Grosshirnrinde in ihren Prinzipien dargestellt auf Grund des Zellenbaues. (Barth, 1909).

3 Tzourio-Mazoyer, N. et al. Automated anatomical labeling of activations in SPM using a macroscopic anatomical parcellation of the MNI MRI single-subject brain. Neuroimage 15, 273–289 (2002).

4 McIntosh, A. et al. Network analysis of cortical visual pathways mapped with PET. Journal of Neuroscience 14, 655–666 (1994).

5 Downing, P. E., Jiang, Y., Shuman, M. & Kanwisher, N. A cortical area selective for visual processing of the human body. Science 293, 2470–2473 (2001).

6 Schubotz, R. I., Anwander, A., Knösche, T. R., von Cramon, D. Y. & Tittgemeyer, M. Anatomical and functional parcellation of the human lateral premotor cortex. Neuroimage 50, 396–408 (2010).

7 Craddock, R. C., James, G. A., Holtzheimer, P. E., Hu, X. P. & Mayberg, H. S. A whole brain fMRI atlas generated via spatially constrained spectral clustering. Human brain mapping 33, 1914–1928 (2012).

8 Nieuwenhuys, R., Broere, C. A. & Cerliani, L. Erratum to: A new myeloarchitectonic map of the human neocortex based on data from the Vogt-Vogt school. Brain structure & function 220, 3753–3755, doi:10.1007/s00429-014-0884-8 (2015).

9 Chong, M. et al. Individual parcellation of resting fMRI with a group functional connectivity prior. NeuroImage 156, 87–100 (2017).

10 Blumensath, T. et al. Spatially constrained hierarchical parcellation of the brain with resting-state fMRI. Neuroimage 76, 313–324 (2013).

11 Salehi, M., Karbasi, A., Scheinost, D. & Constable, R. T. in International Conference on Medical Image Computing and Computer-Assisted Intervention. 478–485 (Springer).

12 Smith, S. M. et al. Functional connectomics from resting-state fMRI. Trends in cognitive sciences 17, 666–682, doi:10.1016/j.tics.2013.09.016 (2013).

13 Gordon, E. M. et al. Generation and evaluation of a cortical area parcellation from resting-state correlations. Cerebral cortex 26, 288–303 (2014).

14 Power, J. D. et al. Functional network organization of the human brain. Neuron 72, 665–678, doi:10.1016/j.neuron.2011.09.006 (2011).

15 Shen, X., Tokoglu, F., Papademetris, X. & Constable, R. T. Groupwise whole-brain parcellation from resting-state fMRI data for network node identification. Neuroimage 82, 403–415 (2013).

16 Thomas Yeo, B. T. et al. The organization of the human cerebral cortex estimated by intrinsic functional connectivity. Journal of Neurophysiology 106, 1125–1165, doi:10.1152/jn.00338.2011 (2011).

17 Fan, L. et al. The human brainnetome atlas: a new brain atlas based on connectional architecture. Cerebral cortex 26, 3508–3526 (2016).

18 Glasser, M. F. et al. A multi-modal parcellation of human cerebral cortex. Nature 536, 171–178 (2016).

19 Van Essen, D. C. Cartography and connectomes. Neuron 80, 775–790, doi:10.1016/j.neuron.2013.10.027 (2013).

20 Eickhoff, S. B., Constable, R. T. & Yeo, B. T. Topographic organization of the cerebral cortex and brain cartography. Neuroimage 170, 332–347 (2018).

21 Fox, P. T. & Lancaster, J. L. Mapping context and content: the BrainMap model. Nature Reviews Neuroscience 3, 319 (2002).

22 Yarkoni, T., Poldrack, R. A., Nichols, T. E., Van Essen, D. C. & Wager, T. D. Large-scale automated synthesis of human functional neuroimaging data. Nature methods 8, 665 (2011).

23 Eickhoff, S. B. et al. Co-activation patterns distinguish cortical modules, their connectivity and functional differentiation. Neuroimage 57, 938–949 (2011).

24 Rosenberg, M. D., Finn, E. S., Constable, R. T. & Chun, M. M. Predicting moment-to-moment attentional state. Neuroimage 114, 249–256 (2015).

25 Esterman, M., Noonan, S. K., Rosenberg, M. & DeGutis, J. In the zone or zoning out? Tracking behavioral and neural fluctuations during sustained attention. Cerebral Cortex, bhs261 (2012).

26 Rosenberg, M. D. et al. A neuromarker of sustained attention from whole-brain functional connectivity. Nat Neurosci 19, 165–171, doi:10.1038/nn.4179 http://www.nature.com/neuro/journal/v19/n1/abs/nn.4179.html#supplementary-information (2016).

27 Rosenberg, M., Noonan, S., DeGutis, J. & Esterman, M. Sustaining visual attention in the face of distraction: A novel gradual-onset continuous performance task. Attention, Perception, & Psychophysics 75, 426–439 (2013).

28 Verbruggen, F., Logan, G. D. & Stevens, M. A. STOP-IT: Windows executable software for the stop-signal paradigm. Behavior research methods 40, 479–483 (2008).

29 Delgado, M. R., Nystrom, L. E., Fissell, C., Noll, D. & Fiez, J. A. Tracking the hemodynamic responses to reward and punishment in the striatum. Journal of neurophysiology 84, 3072–3077 (2000).

30 Baron-Cohen, S., Jolliffe, T., Mortimore, C. & Robertson, M. Another advanced test of theory of mind: Evidence from very high functioning adults with autism or Asperger syndrome. Journal of Child psychology and Psychiatry 38, 813–822 (1997).

31 Bijsterbosch, J. D. et al. The relationship between spatial configuration and functional connectivity of brain regions. Elife 7, e32992 (2018).

32 Gordon, E. M. et al. Precision Functional Mapping of Individual Human Brains. Neuron 95, 791–807.e797, doi:10.1016/j.neuron.2017.07.011 (2017).

33 Van Essen, D. C. et al. The WU-Minn Human Connectome Project: an overview. NeuroImage 80, 62–79, doi:10.1016/j.neuroimage.2013.05.041 (2013).

34 Salehi, M., Karbasi, A., Shen, X., Scheinost, D. & Constable, R. T. An exemplar-based approach to individualized parcellation reveals the need for sex specific functional networks. Neuroimage, Submitted 142 (2017).

35 Wang, D. et al. Parcellating cortical functional networks in individuals. Nature neuroscience 18, 1853 (2015).

36 Bellec, P. et al. Impact of the resolution of brain parcels on connectome-wide association studies in fMRI. Neuroimage 123, 212–228 (2015).

37 Van Essen, D. C., Glasser, M. F., Dierker, D. L., Harwell, J. & Coalson, T. Parcellations and hemispheric asymmetries of human cerebral cortex analyzed on surface-based atlases. Cerebral cortex (New York, N.Y.: 1991) 22, 2241–2262, doi:10.1093/cercor/bhr291 (2012).

38 Nieuwenhuys, R. The myeloarchitectonic studies on the human cerebral cortex of the Vogt-Vogt school, and their significance for the interpretation of functional neuroimaging data. Brain structure & function 218, 303–352, doi:10.1007/s00429-012-0460-z (2013).

39 Greene, A. S., Gao, S., Scheinost, D. & Constable, R. T. Task-induced brain state manipulation improves prediction of individual traits. Nature communications 9, 2807 (2018).

40 Finn, E. S. et al. Functional connectome fingerprinting: identifying individuals using patterns of brain connectivity. Nature neuroscience (2015).

41 Gratton, C. et al. Functional brain networks are dominated by stable group and individual factors, not cognitive or daily variation. Neuron 98, 439–452.e435 (2018).

42 Van Essen, D. C. & Glasser, M. F. Parcellating Cerebral Cortex: How Invasive Animal Studies Inform Noninvasive Mapmaking in Humans. Neuron 99, 640–663, doi:10.1016/j.neuron.2018.07.002 (2018).

43 Haxby, J. V. Multivariate pattern analysis of fMRI: the early beginnings. NeuroImage 62, 852–855, doi:10.1016/j.neuroimage.2012.03.016 (2012).

44 Haxby, J. V., Connolly, A. C. & Guntupalli, J. S. Decoding neural representational spaces using multivariate pattern analysis. Annual review of neuroscience 37, 435–456, doi:10.1146/annurev-neuro-062012-170325(2014).

45 Norman, K. A., Polyn, S. M., Detre, G. J. & Haxby, J. V. Beyond mind-reading: multi-voxel pattern analysis of fMRI data. Trends in cognitive sciences 10, 424–430, doi:10.1016/j.tics.2006.07.005 (2006).

46 Molnár, Z. in Neural circuit development and function in the brain 109–129 (Elsevier, 2013).

47 Mountcastle, V. B. Modality and topographic properties of single neurons of cat’s somatic sensory cortex. Journal of neurophysiology 20, 408–434, doi:10.1152/jn.1957.20.4.408 (1957).

48 Cohen, J. R. The behavioral and cognitive relevance of time-varying, dynamic changes in functional connectivity. NeuroImage 180, 515–525, doi:10.1016/j.neuroimage.2017.09.036 (2018).

49 Gordon, E. M. et al. Precision functional mapping of individual human brains. Neuron 95, 791–807. e797 (2017).

50 Joshi, A. et al. Unified framework for development, deployment and robust testing of neuroimaging algorithms. Neuroinformatics 9, 69–84 (2011).

51 Scheinost, D. et al. Alterations in anatomical covariance in the prematurely born. Cerebral cortex 27, 534–543 (2015).

52 Satterthwaite, T. D. et al. An improved framework for confound regression and filtering for control of motion artifact in the preprocessing of resting-state functional connectivity data. Neuroimage 64, 240–256 (2013).

53 Conley, M. I. et al. The racially diverse affective expression (RADIATE) face stimulus set. Psychiatry research (2018).

54 Cohen, A. O., Conley, M. I., Dellarco, D. V. & Casey, B. in Society for Neuroscience.

55 Barch, D. M. et al. Function in the human connectome: task-fMRI and individual differences in behavior. Neuroimage 80, 169–189 (2013).

56 Speer, M. E., Bhanji, J. P. & Delgado, M. R. Savoring the past: positive memories evoke value representations in the striatum. Neuron 84, 847–856 (2014).

57 May, J. C. et al. Event-related functional magnetic resonance imaging of reward-related brain circuitry in children and adolescents. Biological psychiatry 55, 359–366 (2004).

58 Glass, L. Moire effect from random dots. Nature 223, 578 (1969).

59 Van Essen, D. C. et al. The WU-Minn human connectome project: an overview. Neuroimage 80, 62–79 (2013).

60 Ugurbil, K. et al. Pushing spatial and temporal resolution for functional and diffusion MRI in the Human Connectome Project. Neuroimage 80, 80–104 (2013).

61 Smith, S. M. et al. Resting-state fMRI in the human connectome project. Neuroimage 80, 144–168 (2013).

62 Glasser, M. F. et al. The minimal preprocessing pipelines for the Human Connectome Project. Neuroimage 80, 105–124 (2013).

63 Gomes, R. & Krause, A. in ICML. 391–398.

64 Nemhauser, G. L., Wolsey, L. A. & Fisher, M. L. An analysis of approximations for maximizing submodular set functions—I. Mathematical Programming 14, 265–294 (1978).

65 Minoux, M. in Optimization Techniques 234–243 (Springer, 1978).

66 Lacadie, C. M., Fulbright, R. K., Rajeevan, N., Constable, R. T. & Papademetris, X. More accurate Talairach coordinates for neuroimaging using non-linear registration. Neuroimage 42, 717–725 (2008).

67 Thomas Yeo, B. et al. The organization of the human cerebral cortex estimated by intrinsic functional connectivity. Journal of neurophysiology 106, 1125–1165 (2011).

68 Smith, S. M. et al. Advances in functional and structural MR image analysis and implementation as FSL. Neuroimage 23, S208–S219 (2004).

