## Supplementary Materials for "There is no single functional atlas even for a single individual: Parcellation of the human brain is state dependent"

**This PDF file includes:**
Figures S1 to S8
Video S1
Captions for Figures S1 to S8

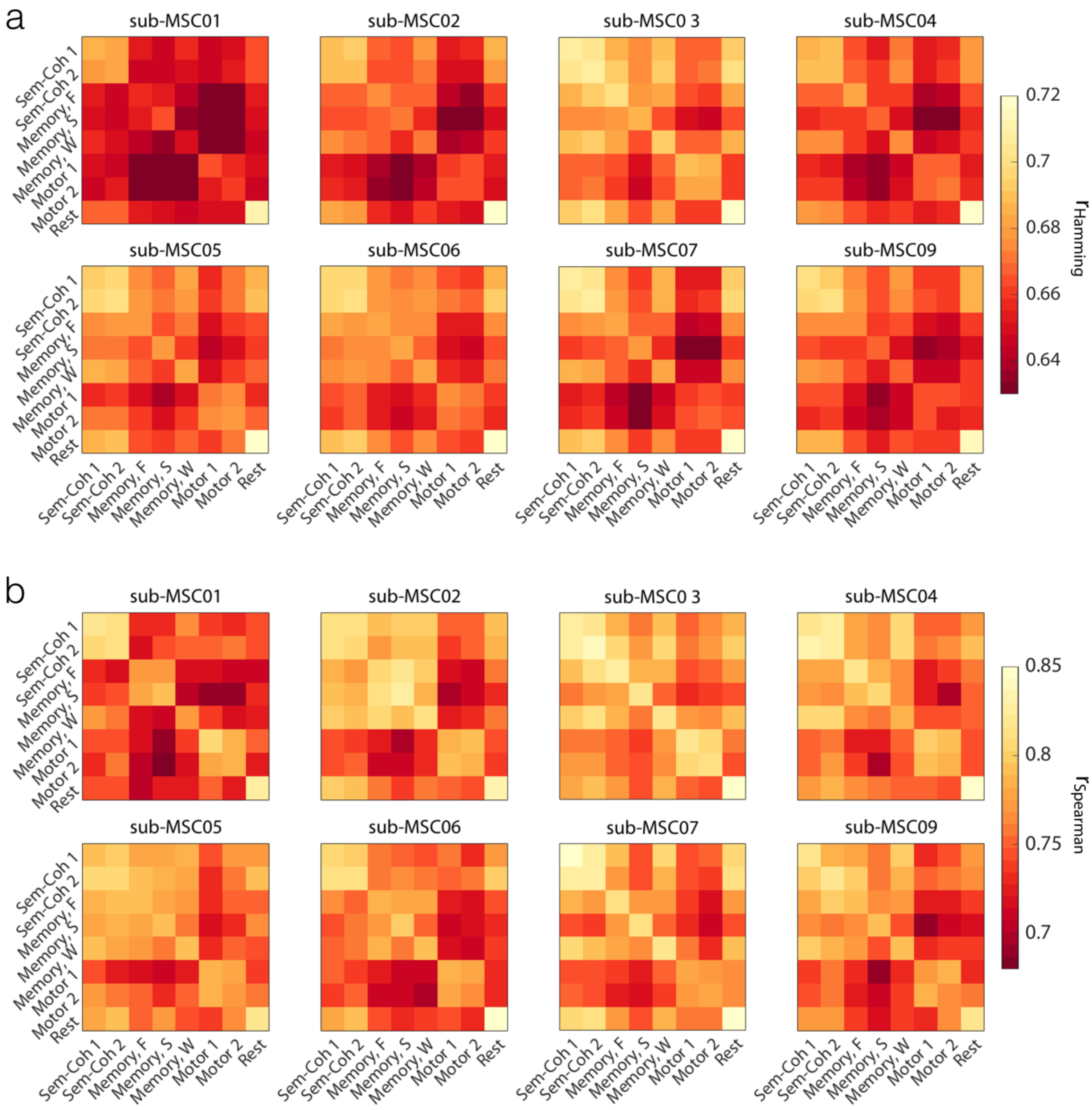


**Figure S1. Individual level replication of the finding that parcel definitions change with task condition; Midnight Scan Club (MSC) data.** Pairwise parcellation similarity was calculated for each individual, within and across functional conditions. For every individual, voting-based ensemble analysis was used with 100 iterations. The matrix represents the average over all iterations. Similarity was assessed by $r_{Hamming} = 1 -$ normalized Hamming distance. b) The same analysis as (a) was performed, this time using rank correlation of parcel-size vectors ($r_{Spearman}$) as a proxy for parcellation similarities.

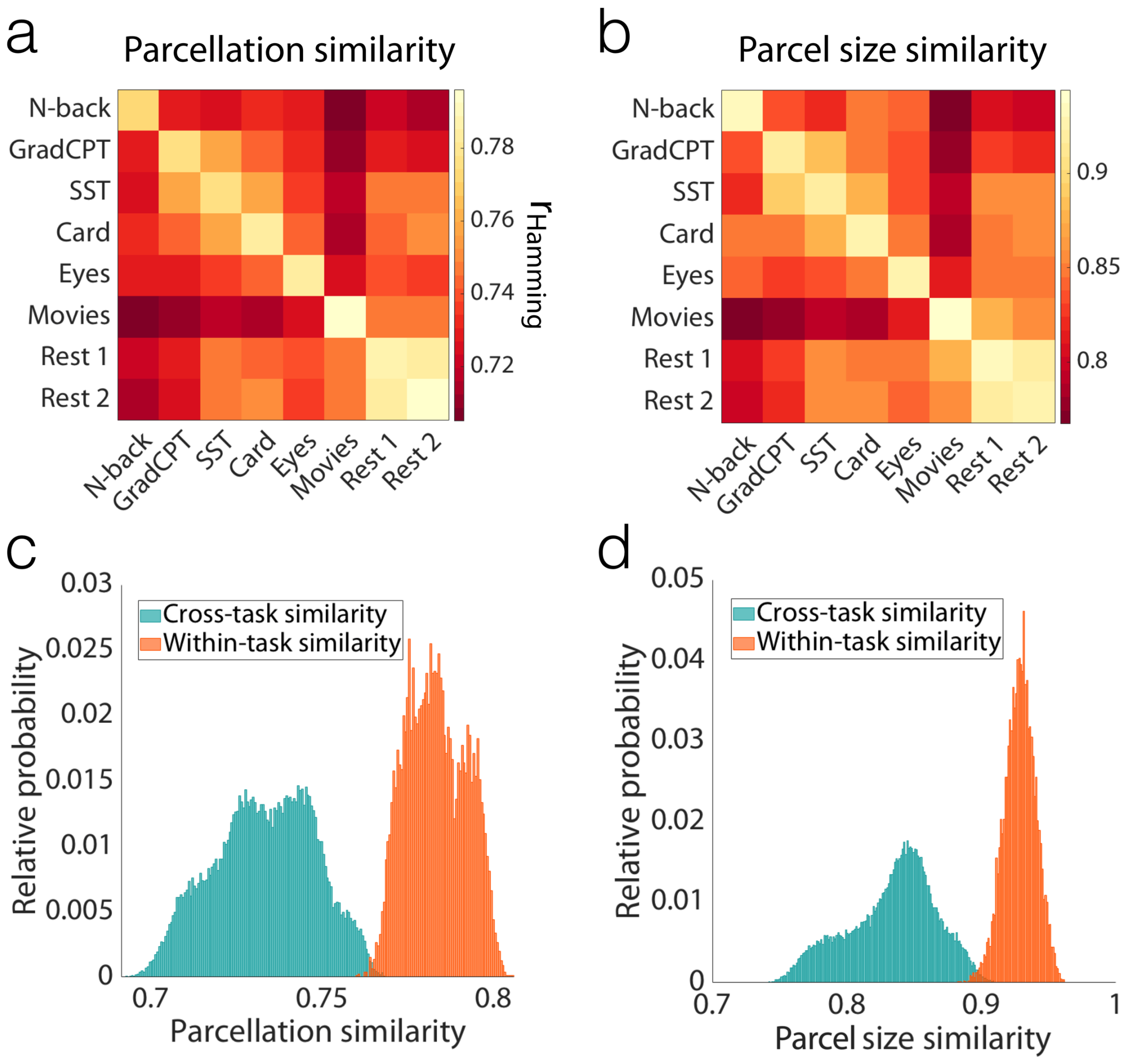


**Figure S2. Replication of the finding that parcel definitions change with task condition, using parcellation of size 368 parcels; Yale data.** a) Pairwise parcellation similarity was calculated within and across functional conditions, using voting-based ensemble analysis with 1000 iterations. Similarity was assessed by $r_{\text{Hamming}} = 1 -$ normalized Hamming distance. b) The same analysis as (a) was performed, this time using rank correlation of parcel-size vectors ($r_{\text{Spearman}}$) as a proxy for parcellation similarities. c) The histogram of the parcellation similarities for all 1000 iterations is depicted for within-condition (diagonal elements in [a]) and cross-condition (off-diagonal elements in [a]) comparisons. The two distributions are significantly different (K-S test; $p<0.001$). d) The histogram of the parcel size similarities for all 1000 iterations is depicted for within-condition (diagonal elements in [b]) and cross-condition (off-diagonal elements in [b]) comparisons. The two distributions are significantly different (K-S test; $p<0.001$).

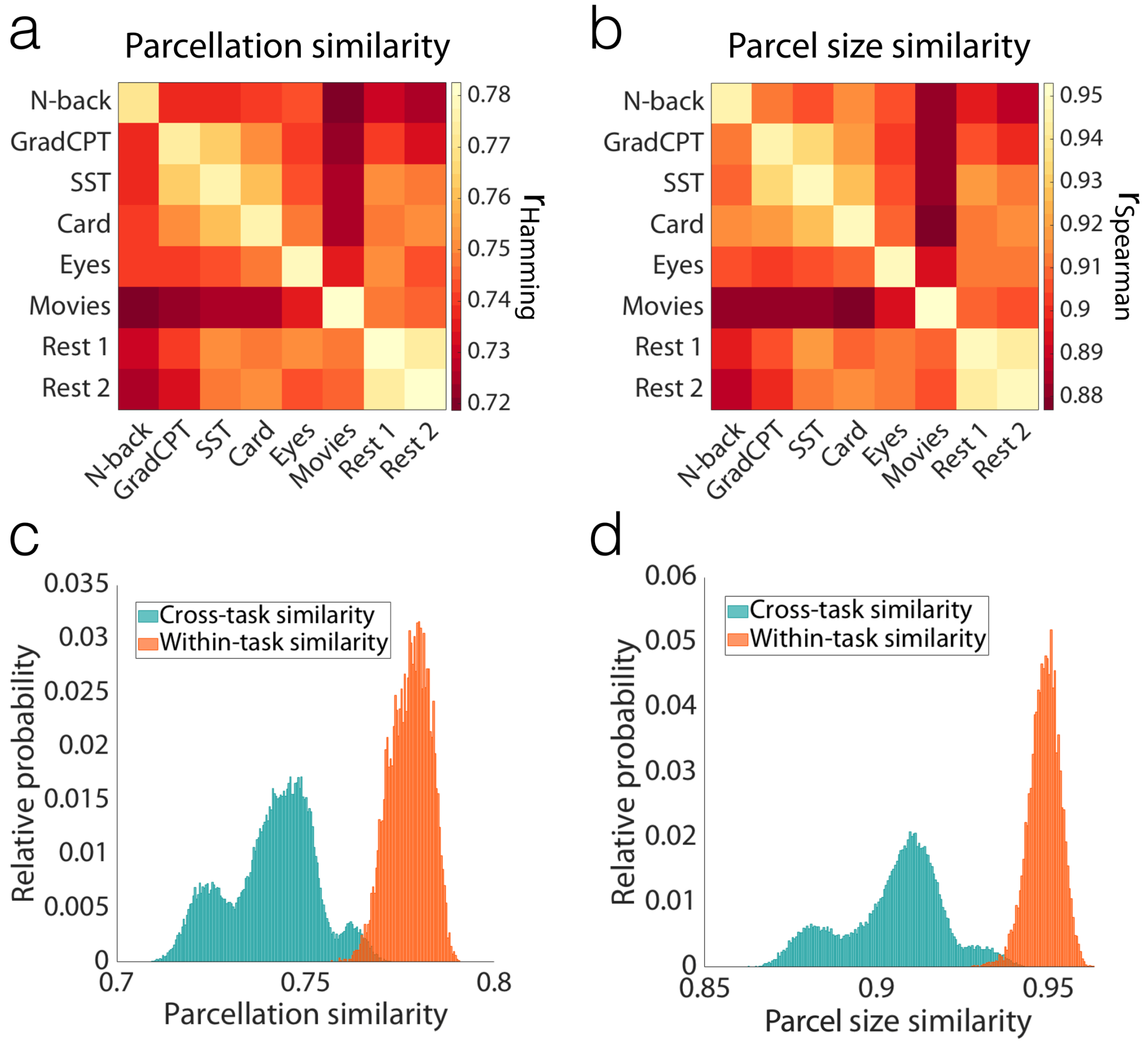


**Figure S3. Replication of the finding that parcel definitions change with task condition, using parcellation of size 1041 parcels; Yale data.** a) Pairwise parcellation similarity was calculated within and across functional conditions, using voting-based ensemble analysis with 1000 iterations. Similarity was assessed by $r_{Hamming} = 1 -$ normalized Hamming distance. b) The same analysis as (a) was performed, this time using rank correlation of parcel-size vectors ($r_{Spearman}$) as a proxy for parcellation similarities. c) The histogram of the parcellation similarities for all 1000 iterations is depicted for within-condition (diagonal elements in [a]) and cross-condition (off-diagonal elements in [a]) comparisons. The two distributions are significantly different (K-S test; $p<0.001$). d) The histogram of the parcel size similarities for all 1000 iterations is depicted for within-condition (diagonal elements in [b]) and cross-condition (off-diagonal elements in [b]) comparisons. The two distributions are significantly different (K-S test; $p<0.001$).

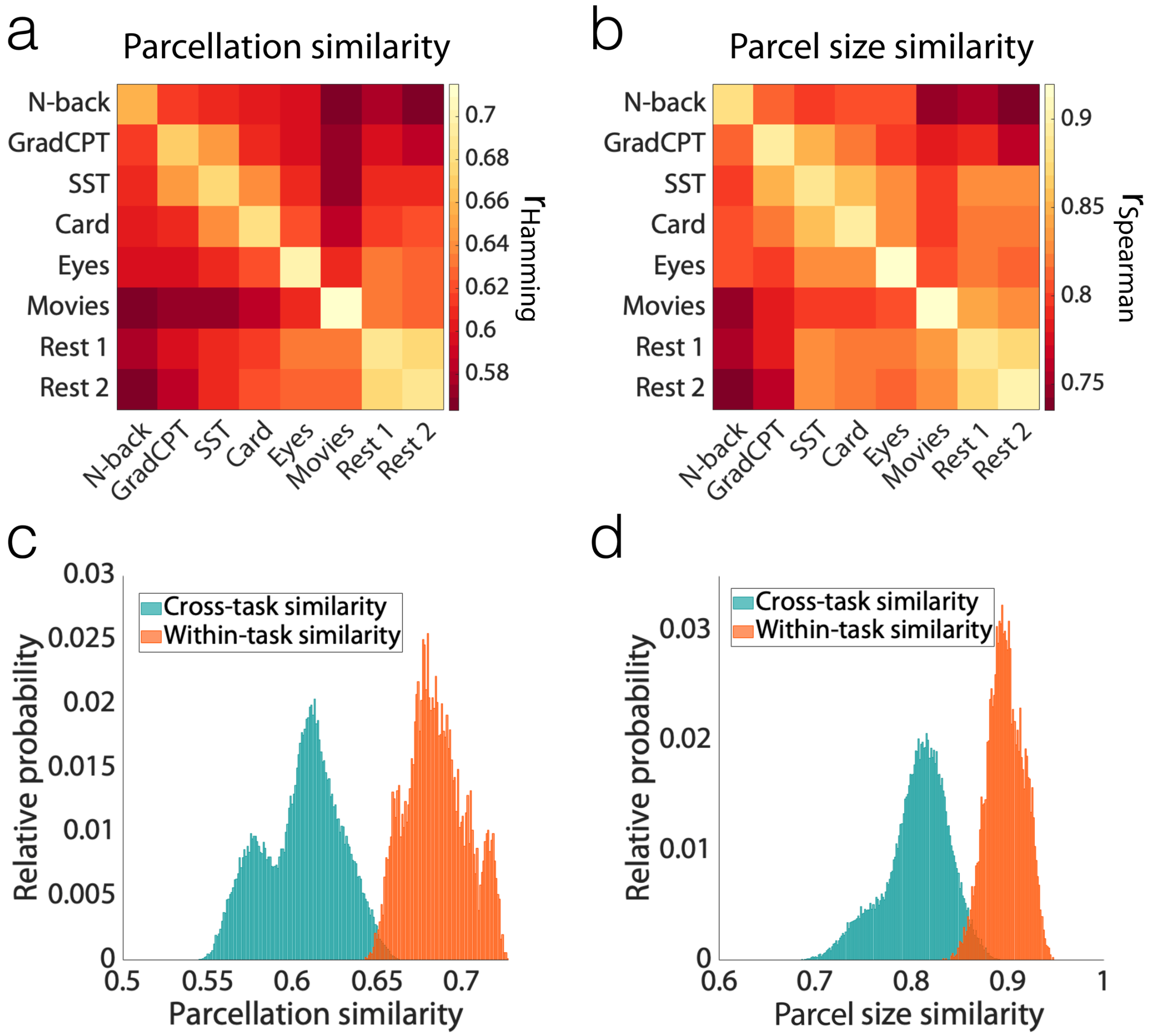


**Figure S4. Replication of the finding that parcel definitions change with task condition, even when exemplar for each parcel is fixed across sessions and conditions; Yale data.** Indication of robustness of the results to the choice of exemplar, such that even with identical exemplars across sessions and conditions, parcels reconfigure reliably based on the brain state. a) Pairwise parcellation similarity was calculated within and across functional conditions, using voting-based ensemble analysis with 1000 iterations. Similarity was assessed by $r_{Hamming} = 1 -$ normalized Hamming distance. b) The same analysis as (a) was performed, this time using rank correlation of parcel-size vectors ($r_{Spearman}$) as a proxy for parcellation similarities. c) The histogram of the parcellation similarities for all 1000 iterations is depicted for within-condition (diagonal elements in [a]) and cross-condition (off-diagonal elements in [a]) comparisons. The two distributions are significantly different (K-S test; $p<0.001$). d) The histogram of the parcel size similarities for all 1000 iterations is depicted for within-condition (diagonal elements in [b]) and cross-condition (off-diagonal elements in [b]) comparisons. The two distributions are significantly different (K-S test; $p<0.001$).

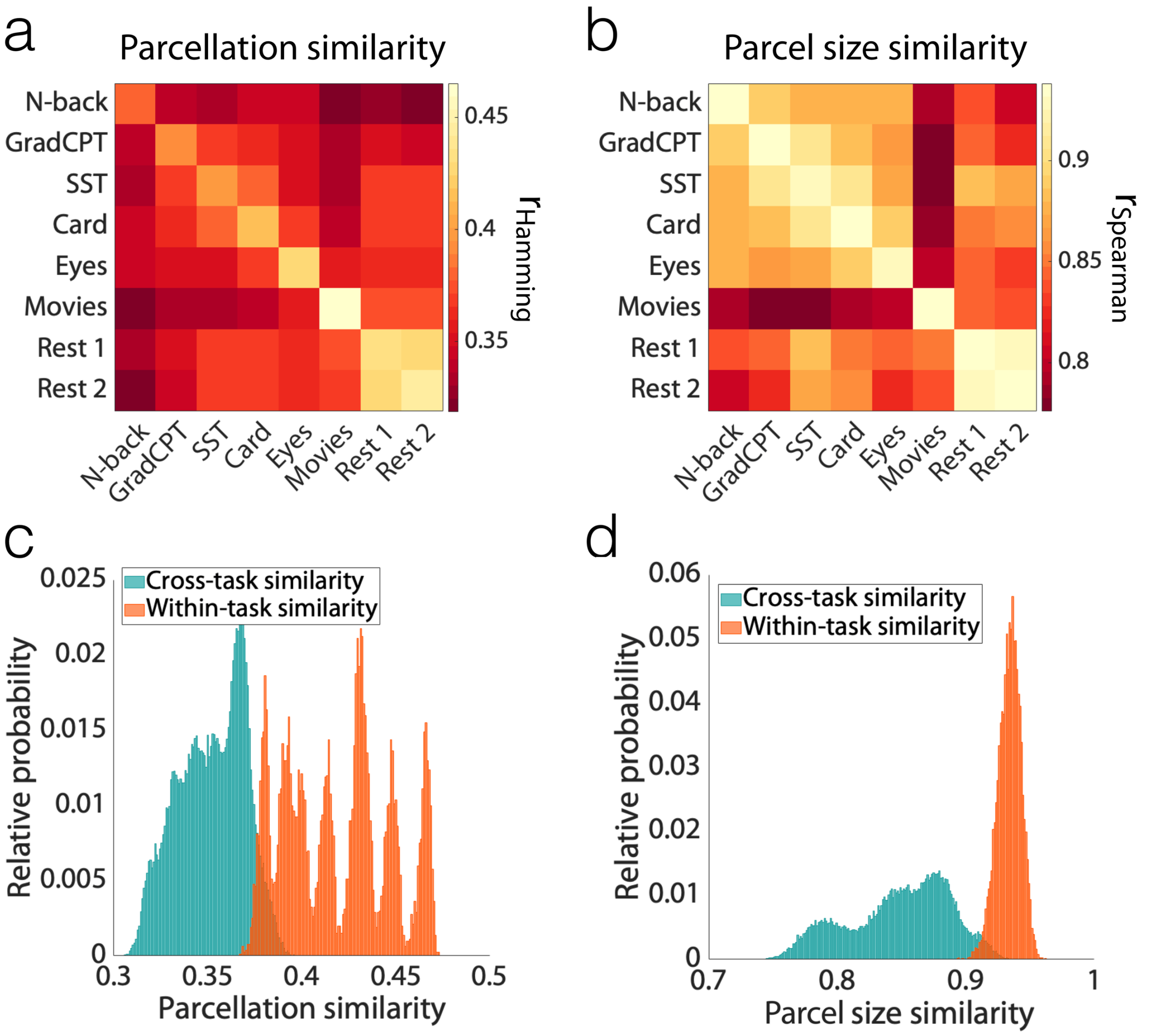


**Figure S5. Replication of the finding that parcel definitions change with task condition, even with a different parcellation algorithm: Wang's iterative parcellation; Yale data.** a) Pairwise parcellation similarity was calculated within and across functional conditions, using voting-based ensemble analysis with 1000 iterations. Similarity was assessed by $r_{Hamming} = 1 -$ normalized Hamming distance. b) The same analysis as (a) was performed, this time using rank correlation of parcel-size vectors ($r_{Spearman}$) as a proxy for parcellation similarities. c) The histogram of the parcellation similarities for all 1000 iterations is depicted for within-condition (diagonal elements in [a]) and cross-condition (off-diagonal elements in [a]) comparisons. The two distributions are significantly different (K-S test; $p<0.001$). d) The histogram of the parcel size similarities for all 1000 iterations is depicted for within-condition (diagonal elements in [b]) and cross-condition (off-diagonal elements in [b]) comparisons. The two distributions are significantly different (K-S test; $p<0.001$).

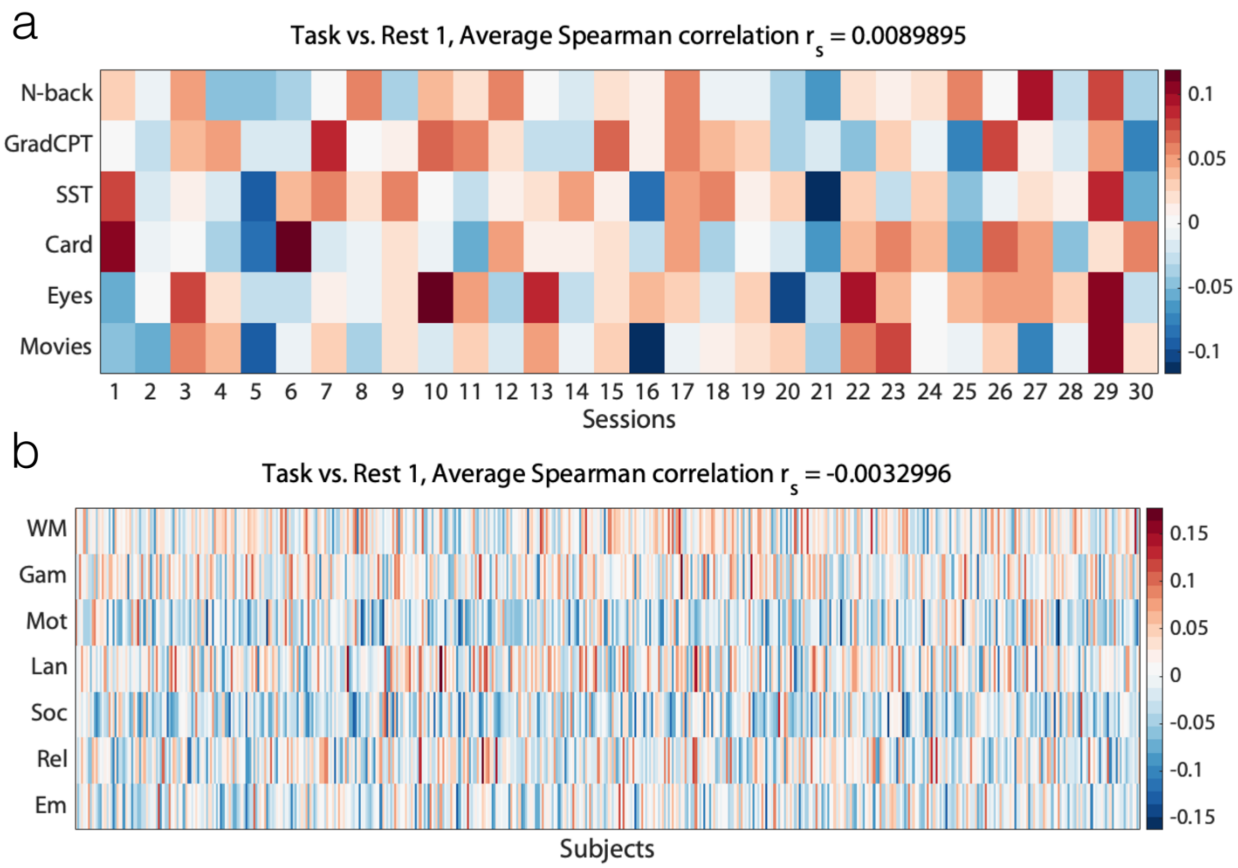


**Figure S6. Correlation between the task activation maps and parcel size differences relative to rest**. The Spearman correlation between the task activation per parcel and the difference in the parcel size (from every task run to the first rest run [Rest 1]) is displayed. (a) Yale data; task activation was approximated for each parcel as the difference between the average temporal signal during each task run and the first rest run (Rest 1). (b) HCP data; task effect size maps were generated using all available individuals' volume-based, FEAT-analyzed, first-level GLM output (COPE files) from the 1200 Subjects Release (S1200) for a given task to generate, using FSL FEAT's FLAME (FMRIB's Local Analysis of Mixed Effects), cross-subject, voxel-wise Cohen's d effect-size contrast maps from t-statistic maps. These values were then averaged for each parcel to yield a mean task effect size per parcel for each task.

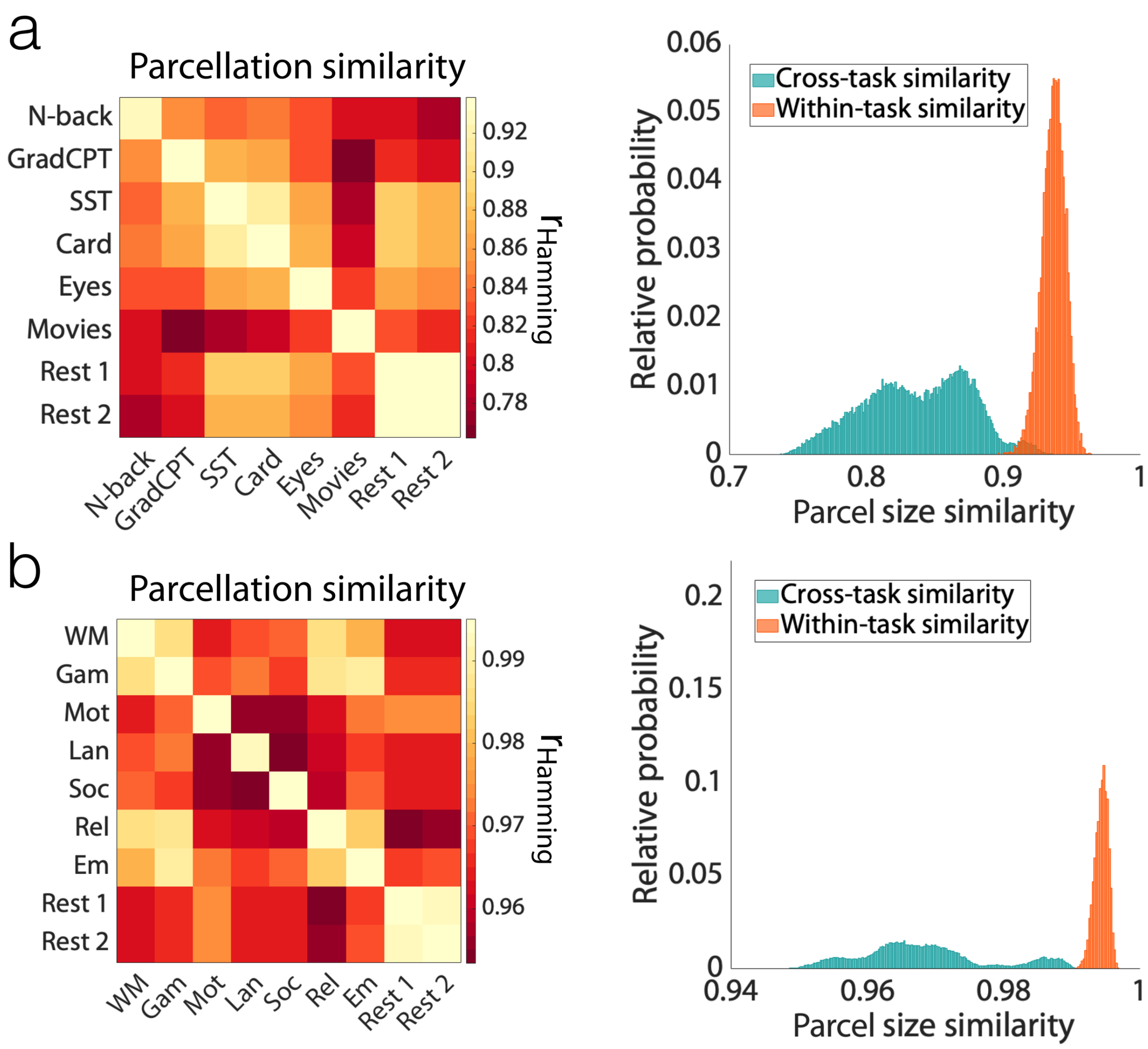


**Figure S7. Replication of the finding that parcel definitions change with task condition, even after eliminating parcels with significant task activation, for (a) Yale data, and (b) HCP data.** Left panels display the pairwise parcellation similarity, calculated within and across functional conditions, using voting-based ensemble analysis with 1000 iterations. Similarity was assessed by rank correlation of parcel-size. Right panels display the histogram of the parcel size similarities for all 1000 iterations for within-condition (diagonal elements in [a]) and cross-condition (off-diagonal elements in [a]) comparisons. The two distributions are significantly different (K-S test; $p<0.001$). a) For Yale data, task activations were approximated per parcel by computing the difference between the average temporal signal during task and rest. b) For HCP data, task effect size maps were generated using all available individuals' volume-based, FEAT-analyzed, first-level GLM output (COPE files) from the 1200 Subjects Release (S1200) for a given task to generate, using FSL FEAT's FLAME (FMRIB's Local Analysis of Mixed Effects), cross-subject, voxel-wise Cohen's d effect-size contrast maps from t-statistic maps. These values were then averaged for each parcel to yield a mean task effect size per parcel for each task.

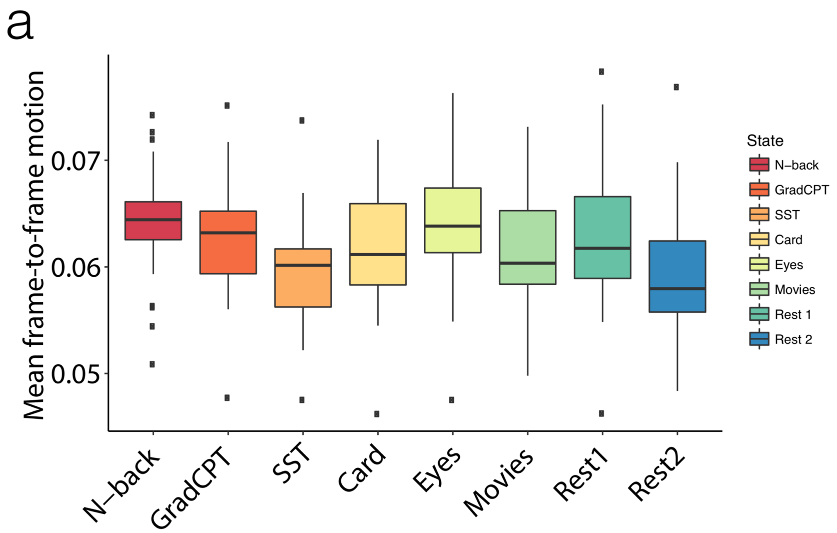


b

| | N-back | GradCPT | SST | Card | Eyes | Movies | Rest 1 |
|---|---|---|---|---|---|---|---|
| **GradCPT** | 1.00 | - | - | - | - | - | - |
| **SST** | 0.02 | 0.31 | - | - | - | - | - |
| **Card** | 1.00 | 1.00 | 1.00 | - | - | - | - |
| **Eyes** | 1.00 | 1.00 | 0.02 | 1.00 | - | - | - |
| **Movies** | 1.00 | 1.00 | 1.00 | 1.00 | 1.00 | - | - |
| **Rest 1** | 1.00 | 1.00 | 1.00 | 1.00 | 1.00 | 1.00 | - |
| **Rest 2** | 0.02 | 0.17 | 1.00 | 1.00 | 0.02 | 0.38 | 0.34 |

**Figure S8. Statistical comparison of head motion across functional conditions; Yale data.** a) Mean frame-to-frame displacement is computed for every functional condition, displayed as a box plot over sessions, where the central mark indicates the median, and the bottom and top edges of the box indicate the 25$^{th}$ and 75$^{th}$ percentiles, respectively. The whiskers extend to the most extreme data points not considered outliers, and the outliers are plotted individually. b) A pairwise Wilcoxon signed-rank test was performed on the mean frame-to-frame displacement. P-values are reported after Bonferroni correction for multiple comparisons.

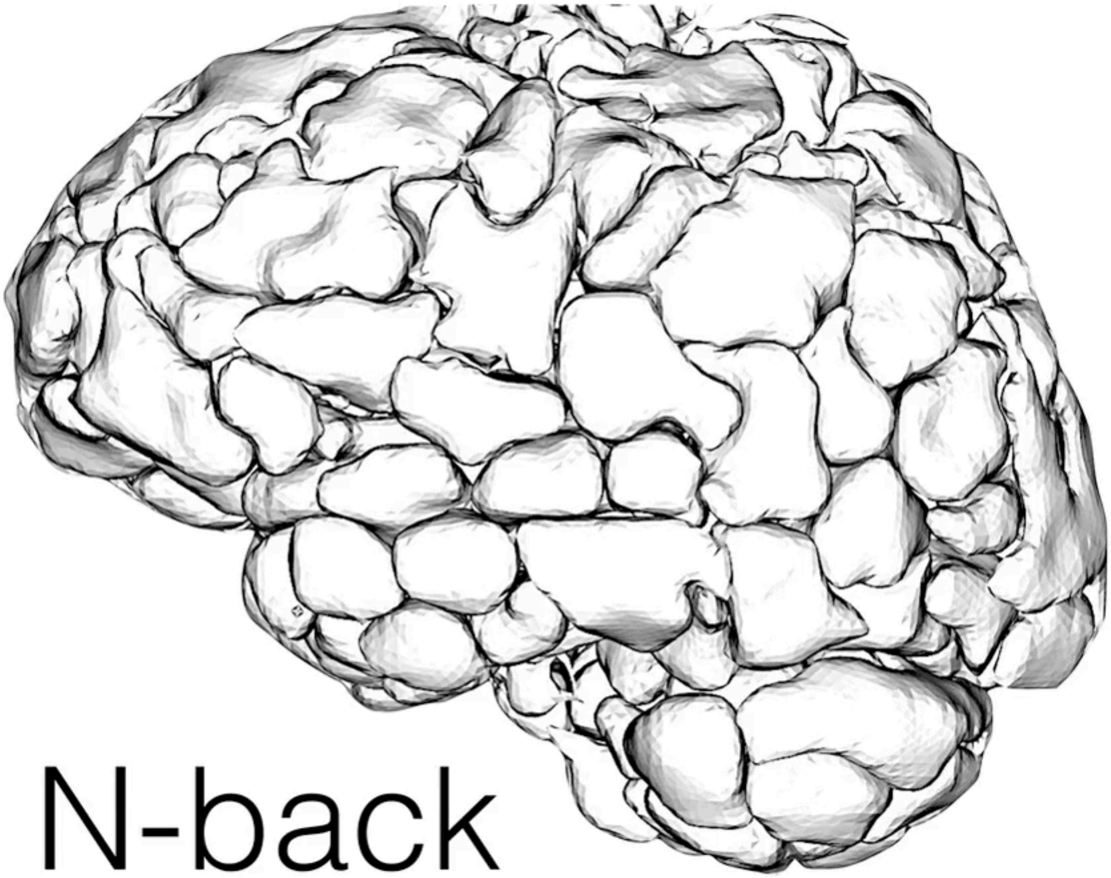


**Video S1. Visualization of the state-specific parcellations; Yale data.**
